# The secreted redox sensor roGFP2-Orp1 reveals oxidative dynamics in the plant apoplast

**DOI:** 10.1101/2025.01.10.632316

**Authors:** Julian Ingelfinger, Lisa Zander, Patricia L. Seitz, Oliver Trentmann, Sophie Tiedemann, Stefanie Sprunck, Thomas Dresselhaus, Andreas J. Meyer, Stefanie J. Müller-Schüssele

**Affiliations:** Molecular Botany, Department of Biology, RPTU Kaiserslautern-Landau, D-67633 Kaiserslautern, Germany; Chemical Signalling, Institute of Crop Science and Resource Conservation (INRES), University of Bonn, D-53117 Bonn, Germany; Cell Biology and Plant Biochemistry, Institute of Plant Sciences, University of Regensburg, D-93053 Regensburg, Germany

**Keywords:** *Arabidopsis thaliana*, *Physcomitrium patens*, redox-sensitive GFP, oxidative burst, genetically encoded biosensor, apoplast, ROS, tip growth

## Abstract

- Specific generation of reactive oxygen species (ROS) is important for signalling and defence in many organisms. In plants, different types of ROS serve useful biological functions in the extracellular space (apoplast), influencing polymer structures as well as signaling during immune responses. The current knowledge of apoplastic ROS dynamics is limited, as dynamic monitoring of extracellular redox processes *in vivo* remains difficult.
- We employed evolutionary distant land plant model species from bryophytes and flowering plants to test whether the genetically encoded redox biosensor roGFP2-Orp1 can be used to assess extracellular redox dynamics.
- Secreted roGFP2-Orp1 can inform about local diffusion barriers and protein cysteinyl oxidation rate in the apoplast, after pre-reduction. Observed re-oxidation rates were slow, within the range of hours. Compared to *Physcomitrium patens*, re-oxidation in *Arabidopsis thaliana* was faster and increased after triggering an immune response. Comparing roGFP2-Orp1 signals in tip-growing *P. patens* protonema and *Nicotiana tabacum* pollen tubes, we consistently find no intracellular redox gradient, but partially reduced extracellular sensor in pollen tubes.
- Our data indicate differences in extracellular oxidative processes between species and within a species, depending on cell type and immune signalling.

## Introduction

Most intracellular compartments, such as the cytosol, plastids and mitochondria constitute a reducing environment for protein cysteinyl groups, as they contain glutathione (GSH) in the mM range as well as a glutathione disulfide reductase that keeps this pool reduced, transferring electrons from NADPH (Marty *et al*., 2009, 2019). In contrast, the extracellular space of plants is characterised by low concentrations of glutathione (less than 1% of total GSH), resulting from GSH or glutathione disulfide (GSSG) export (Ohkama-Ohtsu *et al*., 2007; Zechmann, 2014). GSSG is enzymatically degraded in the apoplast (Ohkama-Ohtsu *et al*., 2007; Noctor *et al*., 2012). Importantly, no thiol-reducing systems are known for the apoplast (Meyer *et al*., 2021). Oxidative protein folding in the secretory pathway prepares proteins for this oxidising environment, ensuring correct formation of disulfides. Measurements of glutathione redox potential (*E*_GSH_) in the ER-lumen using redox-sensitive GFPs (roGFPs) revealed an *E*_GSH_ of less negative than-241 mV (Birk *et al*., 2013; Ugalde *et al*., 2022a).

Although the apoplast seems to be a ‘one-way-street’ in terms of thiol redox biology, redox processes in the apoplast have been implicated in growth, development and reproduction as well as signalling and immunity. To foster cell growth, metabolites constituting building blocks for extracellular polymers, such as the cuticle and cell wall components, are exported (Renault *et al*., 2017). Polymer formation or loosening is induced *in situ*, involving oxidative processes mediated by different ROS, such as superoxide (O_2_^·-^), hydrogen peroxide (H_2_O_2_) and hydroxyl radical formation (Liszkay *et al*., 2004; Tenhaken, 2015; Cosgrove, 2022). Here, class III peroxidases responding with different catalytic cycles to the local H_2_O_2_/O_2_^·-^ balance can mediate formation of hydroxyl radicals that cause cell wall loosening (Chen & Schopfer, 1999; Tenhaken, 2015). Development-and immunity-related cell wall signalling is partially overlapping, often involving membrane-spanning receptor-like kinases (Wolf, 2022). Extracellular O_2_^·-^ and H_2_O_2_ are further involved in evolutionary conserved stress-induced long-distance signalling (Miller *et al*., 2009; Fichman *et al*., 2023; Koselski *et al*., 2023) as well as immune responses. Signal amplification involves the specific formation of O_2_^·-^ via the transmembrane Respiratory Burst Oxidase Homologs (RBOHs), which transfer electrons from the cytosolic NADPH to extracellular oxygen. This triggers a cascade of extracellular ROS formation (Waszczak *et al*., 2018; Mhamdi & Van Breusegem, 2018; Sies *et al*., 2022). After generation, O_2_^·-^ dismutates via enzymatic catalysis or spontaneously at low pH to form the more stable ROS H_2_O_2_ (and O_2_). Oxidative burst signalling and RBOH function in immunity are evolutionary conserved in land plants (Lehtonen *et al*., 2012; Bressendorff *et al*., 2016; Chu *et al*., 2023).

Moreover, extracellular and intracellular redox dynamics can be linked (Mittler *et al*., 2022). Thus, extracellular ROS balance influences the plasma membrane proteome and nanodomain formation (Martinière *et al*., 2019). A first transmembrane redox-responsive receptor kinase (HPCA1) involved in long distance signaling has been described (Castro *et al*., 2021; Fichman *et al*., 2022). Genetically encoded biosensors showed that an extracellular oxidative burst triggered by the application of elicitors is followed by changes of intracellular H_2_O_2_ levels. This apparent correlation suggested that H_2_O_2_ might be transported from the apoplast into the cytosol mediated via peroxiporins and/or diffusion (Nietzel *et al*., 2019; Ugalde *et al*., 2022b; Arnaud *et al*., 2023). The contribution of the cytosolic thiol-based redox regulatory network in H_2_O_2_ detoxification and putative decoding has been further investigated using *in vitro* reconstitution of cytosolic redox-active proteins (Vogelsang *et al*., 2024). Biosensors with high sensitivity to H_2_O_2_-dependent oxidation, such as HyPer7 (Pak *et al*., 2020) and roGFP2-Orp1 (Gutscher *et al*., 2009; Nietzel *et al*., 2019), were also employed successfully to monitor compartment-specific changes in redox states over extended time periods *in vivo* (Nietzel *et al*., 2019; Ugalde *et al*., 2021; Niemeyer *et al*., 2021; Arnaud *et al*., 2023; Dopp *et al*., 2023). Fluorescence read-out of roGFP2 offers high robustness to pH changes, as the protonated chromophore still emits fluorescence, in contrast to many other fluorescent proteins (Schwarzländer *et al*., 2008; Müller-Schüssele *et al*., 2021a). Redox-responsiveness of roGFP2 is mediated by a GSH/GRX-dependent thiol switch on the outer face of the GFP β-barrel structure (Hanson *et al*., 2004; Meyer & Dick, 2010) (**Fig. 1A**). Oxidation rates of roGFP2 can be coupled to H_2_O_2_ levels via proteins containing peroxidatic cysteines, such as yeast Orp1 (Oxidant receptor peroxidase 1, (Ma *et al*., 2007)). This type of protein-based redox sensor is oxidised by H_2_O_2_ and reduced via GSH (Gutscher *et al*., 2009; Nietzel *et al*., 2019) (**Fig. 1A**) and can be expressed in a model organism and compartment of choice. In summary, specific local changes in extracellular ROS levels are involved in immunity, signalling and extracellular polymer formation or loosening. However, it is unclear how dynamic extracellular protein cysteinyl redox states can be, and what oxidation rates are reached *in vivo* in different species. Mechanisms affecting the steady state oxidation levels of cysteine thiols in the apoplast await identification. Staining procedures for different forms of ROS support changes in ROS levels but often cannot resolve temporal or compartmentalised dynamics. Here, we investigate whether secreted roGFP2-Orp1 can be used to sense apoplastic redox dynamics, using evolutionary distant land plant model species with different body plans and alternation of generations, the moss *Physcomitrium patens* and the flowering plants *Arabidopsis thaliana* and *Nicotiana tabacum*.

**Figure 1:**
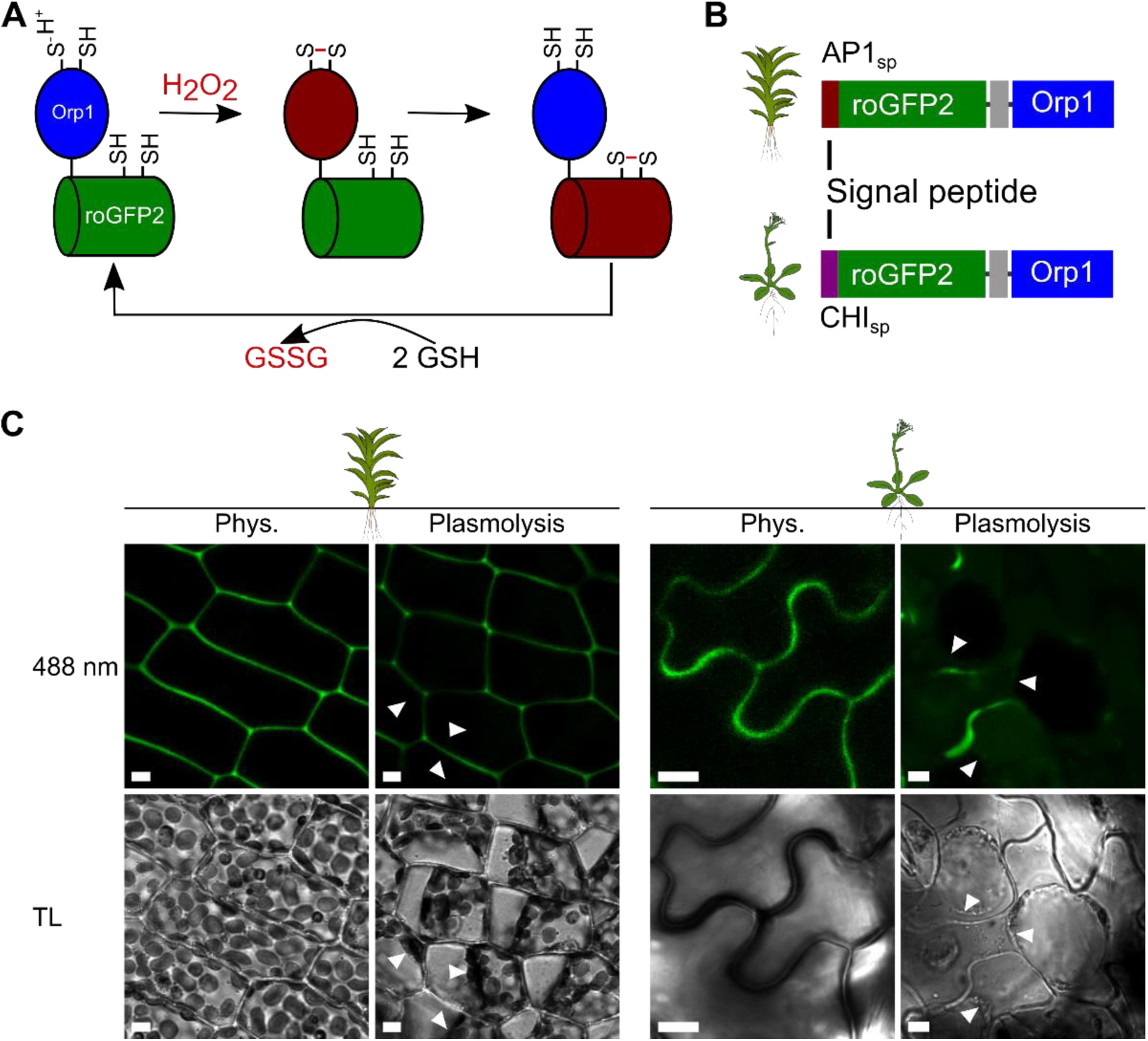
Creation of stable transgenic lines expressing secreted roGFP2-Orp1 in *P. patens* and *A. thaliana*. **A** Simplified scheme showing oxidation and reduction pathways for the redox biosensor roGFP2-Orp1. The reduced form of Orp1 is depicted in blue and for roGFP2 in green, the oxidised forms of both domains are depicted in red. **B** Construct design, adding a suitable signal peptide for secretion to the *roGFP2-linker-Orp1* coding sequence: CHI_sp_ for *A. thaliana* and AP1_sp_ for *P. patens*. **C** Exemplary confocal images of *P. patens* leaflets (line #131) and *A. thaliana* pavement cells (line 3B) stably expressing *AP1_sp_-roGFP2-Orp1* or *CHI_sp_-roGFP2-Orp1*, respectively. Plasmolysis was induced using 0.6 M Mannitol (*A. thaliana*) or 0.6 M NaCl (*P. patens*); arrow heads indicate the plasma membrane. TL, transmitted light; scale bars = 5 µm.

## Material and Methods

### Plant material and growth conditions

*Physcomitrium patens* (Hedw.) Bruch & Schimp ecotype ‘Gransden 2004’ (International Moss Stock Centre (IMSC, http://www.moss-stock-center.org), accession number 40001) was grown axenically and regularly sub-cultured in liquid medium (Knop medium: 250 mg l^-1^ KH_2_PO_4_, 250 mg l^-1^ KCl, 250 mg l^-1^ MgSO_4_ × 7 H_2_O,1 g l^-1^ Ca (NO_3_)_2_ × 4 H_2_O and 12.5 mg l^-1^ FeSO_4_ × 7 H_2_O, pH 5.8) (Reski & Abel, 1985) supplemented with micro-elements (ME), (H_3_BO_3_, MnSO_4_, ZnSO_4_, KI, Na_2_MoO_4_ × 2 H_2_O, CuSO_4_, Co(NO_3_)_2_) (Egener *et al*., 2002). For experimental analyses, *P. patens* gametophores were grown on Knop ME agar plates (12 g l^-1^ purified agar; Oxoid, Thermo Scientific). Light intensity in growth cabinets was set to 70-100 μmol photons m^-2^ s^-1^ and 16:8 h light/dark cycle at 23°C. To induce gametangia development, plates with four-week-old colonies were transferred into a growth cabinet which was set to c. 30 μmol photons m^-2^ s^-1^ and 8:16 h light/dark cycle at 15°C for at least two weeks. *Arabidopsis thaliana* ecotype Columbia 0 (Col-*0*) was used as wild-type (WT) background. Seeds were sown directly on potted soil (ED73 (Patzer Erden,) with 17.6 % (v/v) sand) and stratified in the dark for 48 h at 4°C. Afterwards they were transferred to a plant growth chamber (Fitotron SGC2, Weiss Technik) with light intensity set to c. 130 μmol photons m^-2^ s^-1^ and 8:16 h light/dark cycle at 23°C. After two weeks seedlings were separated with fine forceps, so that only one seedling per pot remained. Four to six weeks after separation the plants were used for experiments.

*Nicotiana tabacum* cv. Petite Havana SR1 plants and *N. benthamiana* were grown in the greenhouse under controlled conditions with a light period of 15h at 28°C and a dark period for 9h at 22°C. Humidity in the chamber was constantly held between 65 and 70%. Anthers from *N. tabacum* were harvested from freshly opened flowers and kept at-80°C until biolistic transformation. Two-week-old *N. benthamiana* seedlings were transplanted into individual pots and used to infiltrate agrobacteria into nearly mature leaves at 4-6 weeks of age.

### Cloning and transient transformation

Both constructs for secreted roGFP2-Orp1 were generated using the GATEWAY™ cloning technology (Invitrogen; Thermo Fisher Scientific). For *P. patens*, the endogenous aspartic protease 1 signal peptide (AP1_sp_, M1-R28 of Pp3c5_19520V3.1) (Schaaf *et al*., 2004) and for *A. thaliana* the endogenous chitinase signal peptide (CHI_sp,_ M1-A23 of AT3G12500.1) (Haseloff *et al*., 1997; Meyer *et al*., 2007; Ugalde *et al*., 2022a) were amplified using PCR (Phusion™ High Fidelity DNA Polymerase, Thermo Fisher Scientific): The AP1_sp_ was amplified, adding an *attB1* site and an overlap to roGFP2 using the primer B1_AP1sp_F GGGGACAAGTTTGTACAAAAAAGCAGGCTTAatgggggcatcgaggagt and AP1sp_ro2Orp_P2 GCCCTTGCTCACCATgcgagggcttgcctcagc. RoGFP2-Orp1 was amplified, adding an *attB2* site and overlap to the AP1_sp_ with the primer AP1sp_ro2Orp_P3 gaggcaagccctcgcATGGTGAGCAAGGGCGAG and B2_ro2_R: GGGGACCACTTTGTACAAGAAAGCTGGGTCTATTCCACCTCTTTCAAAAGTTCT. The fragments were fused in an overlap PCR using the primer B1_AP1sp_F and B2_ro2_R. The resulting AP1_sp_-roGFP2-Orp1 fragment was cloned into *pDONR207* via BP reaction (BP Clonase™II Enzyme Mix, Thermo Fisher Scientific). After sequencing, a positive clone was used for LR reaction (LR Clonase™II Enzyme Mix, Thermo Fisher Scientific) into *PTA2Act5GW* (Bohle *et al*., 2024b), creating a plant expression construct under the control of the *P. patens Actin 5* promoter (Weise *et al*., 2006). To generate the *P. patens* expression vector for cytosolic roGFP2-Orp1, *pENTR207L1L2-roGFP2-Orp1* (Nietzel *et al*., 2019) was used in an LR reaction (LR Clonase™II Enzyme Mix, Thermo Fisher Scientific) into *PTA2Act5GW* (Bohle *et al*., 2024b).

In accordance, the CHI_sp_ was amplified, adding an *attB1* site and an overlap to roGFP2 using the following primer B1_CHIsp_F: GGGGACAAGTTTGTACAAAAAAGCAGGCTTAatgaagactaatctttttctctttc and CHIsp_ro2_P2 CTCGCCCTTGCTCACggcgaattcggccgagga. RoGFP2-Orp1 was amplified, adding an *attB2* site and overlap to the CHI_sp_, using the primer P3_CHIsp_ro2_F: tcggccgaattcgccGTGAGCAAGGGCGAGGAG and B2_ro2(Orp1)_R: GGGGACCACTTTGTACAAGAAAGCTGGGTCTATTCCACCTCTTTCAAAAGTTCT. The two fragments were combined in an overlap PCR using the B1_CHIsp_F and B2_ ro2(Orp1)_R primer. The fragment was cloned into *pDONR207* via BP reaction, generating *pENTR207L1L2-CHI_sp_-roGFP2-Orp1*. After sequencing, a positive clone was used for LR reaction into *pSS02* (derivative of pMDC32 (Curtis & Grossniklaus, 2003)) via LR reaction, creating a plant expression construct under the control of the *A. thaliana Ubiquitin10* promoter.

For transient expression in *N. benthamiana* epidermal leaf cells, the binary expression vector *35S_pro_:CHI_sp_-roGFP2-Orp1* was cloned by Gibson assembly. The *CHI_sp_-roGFP2-Orp1* fragment was amplified from *pENTR207L1L2-CHI_sp_-roGFP2-Orp1* using the primer pair CHIsp_roGFP2_linker_Orp1(F: caatttactattctagtcgaATGAAGACTAATCTTTTTCTCTTTC and R: tgcggactctagcatggccgCTATTCCACCTCTTTCAAAAG). The vector fragment was amplified from the destination vector *pH2GW7* (Karimi *et al*., 2002) with the primer pair pH2GW7 F: CGGCCATGCTAGAGTCCG and R: TCGACTAGAATAGTAAATTGTAATGTTGTTTGTTG.

Both fragments were amplified using KOD Xtreme™ Hot Start DNA Polymerase and ligated via Gibson assembly using NEBuilder® HiFi DNA Assembly Master Mix according to manufacturer’s instructions. Gateway™ LR Clonase™ II was employed to create the binary expression vector *35S_pro_:TagRFP-T-RemA* (plasma membrane marker) by recombining the entry vector *pENTRTagRFP-T-RemA* (Cyprys *et al*., 2019), which contains TagRFP-T N-terminally fused to the membrane anchor of *M. truncatula* SYMREM1 (Konrad *et al*., 2014), with the destination vector *pB2GW7* (Karimi *et al*., 2002). Both expression vectors were transformed into chemically competent *Agrobacterium* strain GV3101 (pMP90RK). Single colonies were picked and grown for 48 h in liquid YEP medium (10 g l^−1^ yeast extract, 10 g l^−1^ peptone, and 5 g l^−1^ NaCl) containing the corresponding antibiotics and cultured to OD_600_ = 0.8 before resuspension in infiltration buffer (5 % (w/v) sucrose, 0,01 % Silwet L-77, 450 µM acetosyringone) supplemented with a small spatula tip of MgSO_4_ and glucose. The two cultures were mixed at 1:1 ratio and co-infiltrated into *N. benthamiana* leaves as described previously for *N. tabacum* (Sparkes *et al*., 2006). After 48 h, plasmolysis was induced by injecting 1 M sorbitol into the infiltrated leaf areas and incubation for 15-30 min before microscopic analysis.

For sensor experiments in *N. tabacum* pollen tubes, the *CHI_sp_-roGFP2-Orp1* and *roGFP2-Orp1* coding sequences were amplified from the *pSS02-CHI_sp_-roGFP2-Orp1* plasmid by Touch Down PCR using KOD Hot Start Polymerase (NOVAGEN) using primers PP373: AACAGGTCTCAGGCTCAATGAAGACTAATCTTTTTCTCTTTCTCAT, PP374: AACAGGTCTCTCTGACTATTCCACCTCTTTCAAAAGTTCT and PP375: AACAGGTCTCAGGCTCAATGGTGAGCAAGGGCG. Amplified DNA fragments and entry vector pGGC000 (Lampropoulos *et al*., 2013) were digested with *BsaI*-HF restriction enzyme (New England Biolabs) to generate compatible overhangs. Ligation into the entry vector was performed with T4 ligase (New England Biolabs). The final expression vectors, driving the constructs under the *LAT52* promotor (Twell *et al*., 1989), were generated by Golden Gate assembly using the Green Gate cloning system and *pGGZ001* as final destination plasmid (Lampropoulos *et al*., 2013). Transient transformation of *N. tabacum* pollen was carried out via particle bombardment, following a protocol adapted from Ge *et al*.(2019). For each transformation, 2 µg of plasmid was mixed with 25 µl of the gold particle (Ø 1.6 µm) suspension. 25 µl of 2.5 M CaCl_2_ and 10 µl of 1 mg ml^-1^ protamine were added to the mixture, vortexed for 3 min and centrifuged at 10, 000 g for 30 s. The resulting pellet was washed with 200 µl of 100% ethanol, vortexed for 3 min, and again centrifuged at 10, 000 g for 30 s before it was resuspended in 16 µl of 100% ethanol. Aliquots of 8 µl each were loaded onto two macro-carriers and air-dried. Pollen was prepared as described by Ge et al. (2019) with the following adaptions: Frozen anthers from 8-10 flowers were suspended in pollen germination medium (PGM) w/o PEG (0.58 mM sucrose, 0.02 M MES KOH, 1.62 µM H_3_BO_3_, 1.66 µM MgSO_4_, 0.98 µM KNO_3_, 3 mM Ca(NO_3_)_2_). After bombardment, pollen grains were transferred into a fresh 6 cm petri dish containing 1.8 ml of PGM with PEG (71.6 µM PEG3350, 73.1 µM sucrose, 0.02 M MES KOH, 1.62 µM H_3_BO_3_, 1.66 µM MgSO_4_, 0.98 µM KNO_3_, 3 mM Ca(NO_3_)_2_) and cultured at 21 °C on a horizontal shaker set to 120 rpm. Germinated pollen grains were either directly transferred onto a microscope slide or treated with 5 mM 2,2’-dipyridyl disulfide (DPS) or 10 mM dithiothreitol (DTT) in PGM for at least 30 min before transfer.

### Generation of transgenic lines

Stable transgenic *P. patens* lines were generated using polyethylene-glycol mediated protoplast transformation, as described in Hohe *et al*. (2004) For transformation, purified plasmids were cut near *PTA2* homologous regions (Kubo *et al*., 2013) (*PTA2Act5GW-AP1sp-roGFP2-Orp1* using *BglII* and *NotI*; *PTA2Act5GW-roGFP2-Orp1* using *BglII)* and mixed in a molar ratio of c. 2:1 with the uncut resistance plasmid *pBsNNNEV*, containing the *nptII* neomycin resistance gene under the control of a *NOS* promoter and terminator. Plants surviving four weeks on selection (Knop ME with 12.5 µg ml^-1^ G418) were transferred to Knop ME and screened for sensor fluorescence after growth for c. 2 weeks.

Transformation of *A. thaliana* was performed by the floral dip method (Clough & Bent, 1998). Briefly, competent *AGL-1 Agrobacterium tumefaciens* cells were transformed with an error-free expression clone of *pSS02-CHI_sp_-roGFP2-Orp1* using electroporation and selected on agar containing Rifampicin, Ampicillin and Kanamycin (all 50 μg ml^-1^). For floral dip, an overnight culture of a positive *A. tumefaciens* colony was inoculated and after reaching an OD_600_ of approximately 0.8, the cells were harvested by centrifugation for 10 min at 5, 000 g. The supernatant was discarded, and the pellet resuspended in 400 ml dipping solution (5% sucrose, 0.02% Silwet Gold). *A. thaliana* plants containing flower buds were dipped twice with a time interval of 6-7 days. T1-transformants were selected on ½ MS (Murashige & Skoog + Vitamins 0.22%, Duchefa), 0.1% sucrose, 0.8% agar plates, pH 5.7, containing 20 µg ml^-1^ Hygromycin B (Duchefa). After germination, seedlings with long hypocotyls, longer roots as wells as green cotyledons were transferred to soil. T1 plants were screened for fluorescence and presence of a single T-DNA was confirmed using segregation analysis of T2 seedlings. Experiments shown were conducted with material from homozygous T4 and T5 plants.

### Microscopic imaging of roGFP2-Orp1

For ratiometric analysis, confocal laser scanning microscopy of roGFP2-Orp1 in the apoplast of *P. patens* leaflets and *A. thaliana* leaf discs was realised on a LSM880 (Axio Observer.Z1, Carl Zeiss) using a x40 (C-Apochromat x40/1.2 W) objective by exciting roGFP2 sequentially at 405 nm (diode laser, 1/0.5 % (*A. thaliana* / *P. patens*)) and 488 nm (argon laser, 1/3.5 % (*A. thaliana* / *P. patens*)) and detecting roGFP2 emission between 509 nm and 535 nm. Autofluorescence was detected at 425-470 nm after excitation at 405 nm. Chlorophyll autofluorescence was detected at 680-735 nm after excitation at 488 nm. For roGFP2-Orp1 calibration, *P. patens* gametophores and *A. thaliana* leaf discs were submerged in imaging buffer (10 mM MES, 5 mM KCl, 10 mM CaCl_2_, 10 mM MgCl_2_ pH 5.8 (Wagner *et al*., 2019)) or imaging buffer supplemented with either 5 mM 2,2’-dipyridyl disulfide (DPS) or 10 mM dithiothreitol (DTT) and incubated until complete oxidation or reduction. Fluorescence ratio calculations and further image analysis was performed in a MATLAB-based ratio software (RRA) (Fricker, 2016). For plasmolysis, 600 mM Mannitol was used for *A. thaliana* and 600 mM NaCl was used for *P. patens*. The degree of oxidation (OxD) of roGFP2 was calculated according to Schwarzländer *et al*. (2008).

Overview images of fluorescence intensity were taken at a fluorescence stereomicroscope (Leica M205 FCA with Leica K3C camera, Plan Apo 0.63x and 2x corr. objective), using 405-40 ex./525-50 em. and 470-40 ex./525-50 em. filter combinations.

Transiently transformed *Nicotiana benthamiana* leaves were imaged using a Visitron VisiScope spinning disc microscope with a Leica DMi8 body, a Yokogawa CSU-W1 spinning disc unit, a Visitron sCMOS pco.edge camera, and a 40x/1.2 water immersion objective. The 488 nm laser was used for excitation of GFP, tagRFP was excited with the 561 nm laser. Emission filters were 525-50 nm for GFP and 609-52 nm for RFP. Images were processed using Fiji (Schindelin *et al*., 2012).

Fluorescence microscopy of *N. tabacum* pollen tubes was performed using a ZEISS LSM 980 Airyscan2 Confocal Laser Scanning Microscope (CLSM) with a Plan-Apochromat 63x/1.40 Oil DIC M27 objective. CHI_sp_-roGFP2-Orp1/roGFP2-Orp1 were excited sequentially with a 405 nm (8%) and 488 nm (4%) laser and detected at 499-535 nm. Autofluorescence was detected for both sensor constructs at 400-477 nm after excitation at 405 nm.

### Plate reader-based read-out of fluorescence and luminescence

Ratiometric time-series measurements for roGFP2-Orp1 fluorescence were carried out using a CLARIOstar® Plus plate reader (BMG Labtech). The roGFP2 signal was detected using a sequential filter-based excitation of 400-10 nm and 482-16 nm, with the emission detected using a 530-40 nm filter, using top optics. For *P. patens* expressing AP1_sp-_roGFP2-Orp1, whole gametophores were transferred to a 96-well plate (3-5 per well) and incubated in 200 µl imaging buffer (10 mM MES, 5 mM KCl, 10 mM CaCl_2_, 10 mM MgCl_2_ pH 5.8) overnight in the dark at room temperature. The next day, the imaging buffer was replaced and, after another hour in the dark, the measurement was started (Ugalde *et al*., 2022a). After measuring the initial fluorescence of the untreated samples, the measurement was paused and the treatment was added. For calibration the treatment of 10 mM DTT for full reduction and 5 mM DPS for full oxidation was used, while for re-oxidation measurements the reductive treatment consisted of 5 mM tris(2-carboxyethyl)phosphine (TCEP) (Bond-Breaker™ TCEP-solution, REF 77720, Thermo Scientific). In this case, the measurement was paused again to remove and wash out the TCEP 3 x 5 min with 200 µl imaging buffer, after observing complete sensor reduction.

For assays with *A. thaliana* expressing *CHI_sp_-roGFP2-Orp1*, 7 mm leaf discs were punched out and floated on imaging buffer overnight in the dark with the adaxial side facing up. On the next day, the leaf discs were transferred to a 96-well plate and pushed to the bottom of the well with the adaxial side facing down. After one hour of incubation in imaging buffer in the dark, the buffer was replaced and leaf discs used for fluorescence read-out of roGFP2-Orp1 as described above, but via bottom optics (Ugalde *et al*., 2022a). To trigger oxidative bursts, a flagellin fragment (*flg22*, AS-62633, Eurogentec) was used as elicitor and added manually to a final concentration of 1 µM in the respective wells.

For a luminescence-based read-out of elicitor triggered oxidative bursts (Luminol assay), 4 mm leaf disks of *A. thaliana* Col-*0* were prepared and incubated overnight as described above. They were then transferred to imaging buffer or 5 mM TCEP in imaging buffer for 2 hours and subsequently washed 3 x 5 min with imaging buffer. Afterwards, they were transferred to a 96-well plate and left floating on imaging buffer, containing the assay mixture of 20 µM L-012 (L-012, 120-04891, FUJIFILM Wako Chemicals Europe GmbH) and 8 µg/ml horseradish peroxidase (HRP1) (P8125, Sigma-Aldrich). The relative light units (RLU) were measured for at least 5 cycles before an oxidative burst was triggered as described above. As a control, imaging buffer was added instead of *flg22*. To allow for comparison between replicates on different plates, RLU data of each plate were normalised over the mean peak RLU value of control leaf discs treated with *flg22*.

## Statistical analysis

Two-way ANOVA followed by Tukey’s multiple comparisons test was performed using GraphPad Prism version 10.4.0 (or later versions) for Windows (GraphPad Software, Boston, Massachusetts USA, www.graphpad.com).

## Results

### Generation of stable plant lines secreting roGFP2-Orp1 to the apoplast

To obtain stable plant lines expressing and secreting roGFP2-Orp1 (**Fig. 1A**) to the apoplast, we used endogenous constitutive promoters and added suitable signal peptides for targeting to the secretory pathway to the N-terminus of existing roGFP2-Orp1 constructs (Nietzel *et al*., 2019). To this end, we employed the endogenous signal peptide of aspartic protease 1 (AP1_sp_) from *P. patens* (Schaaf *et al*., 2004; Hoernstein *et al*., 2018), as well as an endogenous chitinase signal peptide from *A. thaliana* (Haseloff *et al*., 1997; Meyer *et al*., 2007; Ugalde *et al*., 2022a) (**Fig. 1B**). Correct protein targeting to the apoplast was confirmed by transient expression of *CHI_sp_-roGFP2-Orp1* together with a plasma membrane marker in *Nicotiana benthamiana* epidermal cells (**Fig. S1**).

Subsequently, we stably transformed *P. patens* and *A. thaliana* and selected several independent fluorescent plant lines expressing secreted roGFP2-Orp1. No growth differences were observed for transgenic lines and two independent lines per species were chosen for further analyses. By inducing plasmolysis, we confirmed the localisation of fluorescent signal in the apoplast for both species (**Fig. 1C, Fig. S2**). While roGFP2 fluorescence was evenly distributed in leaflets of moss gametophores (**Fig. 1C**, **Fig. S2**, **Fig. S3A, B**), it was mainly visible in perpendicular walls in filamentous moss protonema (**Fig. S3C**). In *A. thaliana*, roGFP2 fluorescence signal intensities differed between tissues: the strongest signals were observed in lobes of pavement cells in epidermis as well as hypocotyls of seedlings (**Fig. 1C, Fig. S3D, E, F**), whereas mesophyll cells showed a heterogenous signal distribution with punctuate signals at contact points between cells (**Fig. S3G**). Sensor fluorescence intensity in roots was lower, except for the root tip (**Fig. S3H, I**).

### Calibration of secreted roGFP2-Orp1

Next, we assessed the redox state of secreted roGFP2-Orp1 by ratiometric imaging and ratiometric plate reader-based fluorescence read-out for both species. To test functionality and dynamic range of secreted roGFP2-Orp1, we fully reduced and oxidised the sensor protein *in vivo*, using the reductant dithiothreitol (DTT) and the thiol-specific oxidant 2,2’-dipyridyl disulfide (DPS) (Lopez-Mirabal *et al*., 2007). Using confocal microscopy and image analysis (Fricker, 2016; Ugalde *et al*., 2022a), we found that roGFP2-Orp1 dynamic range (δ) was between c. 1.8 and 2.5 in moss leaflets (**Fig. 2A, B** individual channels shown in **Figs. S4**), while it was increased in the reproductive structures, the gametangia, to c. 3 to 4 (**Fig. S5, S6**). We found a similar dynamic range of c. 1.8 to 2.3 in *A. thaliana* leaf epidermis and mesophyll (**Fig. 2A, B**; individual channels shown in **Fig. S7, S8**). Using plate reader-based fluorescence read-out from gametophores and leaf discs (Wagner *et al*., 2019; Ugalde *et al*., 2022a), we measured a slightly smaller or similar dynamic range between c. 1.8 and 2.3 during sensor calibration over time in both species (**Fig. 2C**). As calibration was possible in all tested tissues, it is possible to determine the *in vivo* redox state of roGFP2-Orp1 by investigating non-treated, physiological samples (phys.). Both ratiometric measurement methods confirmed that roGFP2-Orp1 is mainly oxidised when secreted (data for physiological state without treatments in **Fig. 2A, B, C**), with many physiological values not significantly different from fully oxidised sensor controls. Based on the individual calibrations, we calculated mean degrees of sensor oxidation (OxD) in physiological state ranging from 65%+/-13% to 87%+/-5% for *P. patens* and 75% +/-3% to 90% +/-6% in *A. thaliana*, which is similar to the 96.5 % that was shown in Arnaud *et al*. (2023).

**Figure 2:**
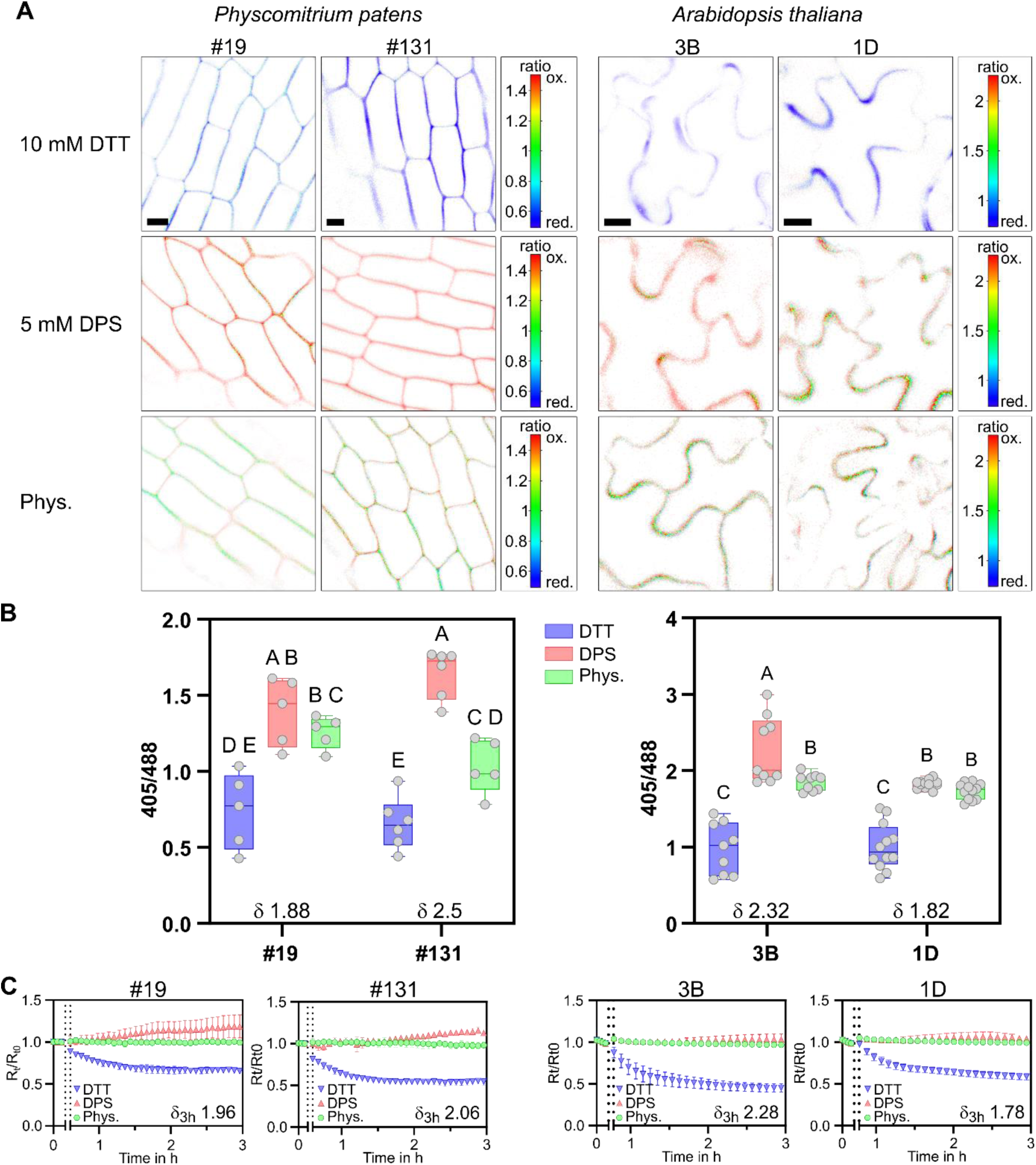
**Calibration of apoplastic roGFP2-Orp1 in *P. patens* and *A. thaliana*** Calibration is shown for two independent lines of *P. patens* expressing *AP1_sp_-roGFP2-Orp1* (left panels) and two independent lines of *A. thaliana* expressing *CHI_sp_-roGFP2-Orp1* (right panels). 10 mM dithiothreitol (DTT) was used for full reduction and 5 mM 2,2’-dipyridyl disulfide (DPS) was used for full oxidation of the biosensor *in vivo*. **A** Sensor 405/488 redox ratio depicted as false-colour ratio scale in confocal microscopy images. Scale bars = 10 µm. **B** Box plots of redox ratio analysis using confocal microscopy images. The measured dynamic range is indicated by δ. Boxes show 25th to 75th percentiles, the line indicates the median, the whiskers show min and max, individual data points are shown as circles, n = 5-12. Different letters indicate significant differences according to a two-way ANOVA with Tukey’s multiple comparison post-hoc test, p < 0.05. **C** *In vivo* calibration of roGFP2-Orp1 measured as a time series using leaf discs (*A. thaliana*) or gametophores (*P. patens*) in a multi-well plate reader. Graphs depict 400-10/482-16 ratios, normalised to the ratio mean before treatment (R_t_/R_t0_). Opening of the plate reader for addition of DTT or DPS, respectively, and start of the treatment is indicated by two dashed lines, n = 5.

Plate reader-based measurements, that integrate biosensor signal over a high number of cells, suggested that secreted roGFP2-Orp1 was not completely oxidised *in vivo* in *P. patens*, as plateau mean values for physiological measurements were consistently lower than mean values for oxidised controls (**Fig. 2C**). Interestingly, using confocal microscopy, some heterogeneity of sensor oxidation was visible both in *A. thaliana* and *P. patens*, especially in areas of high sensor signal **(Fig. 2 A&B, Fig. S4, Fig. S7, Fig. S8).**

### Monitoring protein cysteinyl oxidation kinetics in the apoplast

While the oxidation rate of roGFP2-Orp1 is dependent on changes in H_2_O_2_ levels, reduction of the roGFP2 reporter domain is dependent on the GSH/GRX system intracellularly. Given that GRXs are absent from the apoplast and that GSH levels are very low, the sensor oxidation should be uncoupled from competing reductive processes. Therefore, the sensor oxidation state could potentially be used as a read-out for changes in apoplastic H_2_O_2_ levels, if the sensor disulfides can be pre-reduced. Thus, we tested different doses and types of thiol-reducing agents, using the stable lines expressing secreted roGFP2-Orp1. To this end, we first used DTT, that already proofed successful in apoplastic sensor calibration (**Fig. 2**). After reaching plateau reduction levels, we subsequently removed DTT, washed the samples stringently and investigated sensor re-oxidation rates (**Fig. S9**). A direct comparison of *P. patens* and *A. thaliana*, including sensor calibration, revealed that secreted roGFP2-Orp1 oxidation rates after pre-reduction showed species-specific differences, with barely any re-oxidation observable for *P. patens* gametophores and re-oxidation in *A. thaliana* leaf discs within the range of hours (**Fig. S9**). To titrate exposure to the reducing agent DTT, we reduced DTT doses from 10 mM to 1 mM and found that 2.5 mM DTT was sufficient to fully reduce secreted roGFP2-Orp1 in approximately 2-4 h in both species (**Fig. S10**). Re-oxidation occurred only slowly in *A. thaliana* leaf discs in a period of c. 10 h and using 5 mM or more DTT led to incomplete sensor re-oxidation in the monitored time period of c. 20 h. Notably, re-oxidation in *P. patens* remained consistently low, independently of the used amount of DTT **(Fig. S9)**. As DTT penetrates cells and as complete reduction of cells might cause reductive stress and disturb ER function (Ugalde *et al*., 2022a), we next tested a reducing agent that is non-cell permeable. To this end, we used 5 mM TCEP (Cline *et al*., 2004). As a control, we tested TCEP effects using *A. thaliana* leaf discs expressing cytosolic roGFP2-Orp1 and found no effect during 4 h incubation (**Fig. S11**). Interestingly, cytosolic roGFP2-Orp1 showed a potential oxidation after removal of TCEP (**Fig. S11**). Incubating *A. thaliana* leaf discs with 5 mM TCEP led to complete reduction of secreted roGFP2-Orp1, while full dynamic range was not reached in *P. patens* gametophores (**Fig. 3A**). After washing out TCEP, secreted roGFP2-Orp1 re-oxidised in a range of c. 12 to 30 h in independent experiments in *A. thaliana*, while its redox state remained largely unchanged in *P. patens* gametophores. To test if roGFP2-Orp1 was still responsive to H_2_O_2_-mediated oxidation after pre-reduction, we added exogenous H_2_O_2_ after removal of reducing agent and observed rapid oxidation responses of secreted roGFP2-Orp1 *in vivo* in both species (**Fig. 3B**).

**Figure 3:**
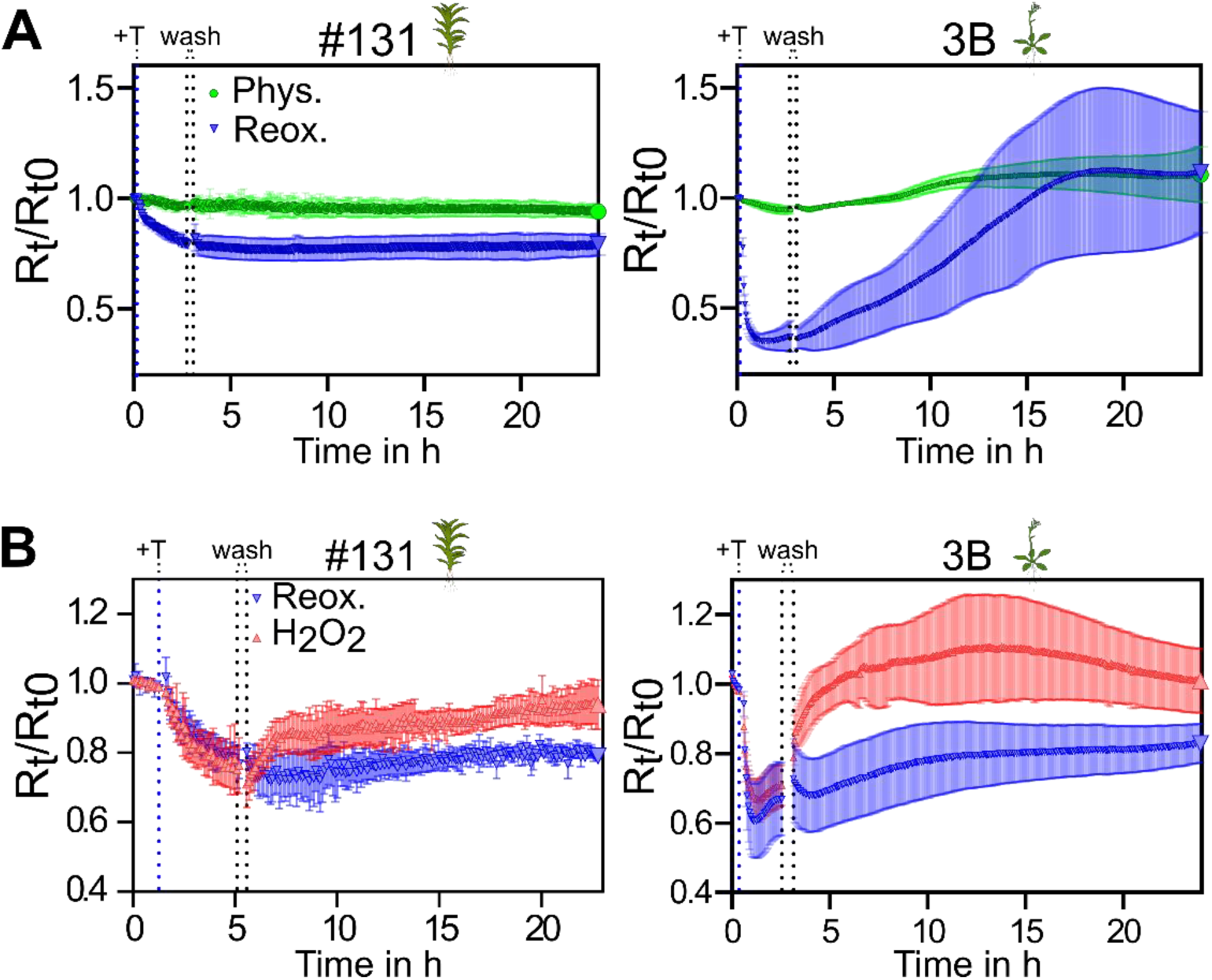
Re-oxidation rate of reduced apoplastic roGFP2-Orp1. Graphs depict 400-10/482-16 ratios, normalised to the ratio mean before treatment (R_t_/R_t0_), of apoplastic roGFP2-Orp1 measured as a time series using leaf discs (*A. thaliana*) or gametophores (*P. patens*) in a multi-well plate reader. Start of treatment with 5 mM TCEP is indicated by a blue dashed line (+T). After reaching plateau values the measurement was paused to remove and wash out the reducing agent, indicated by the two dashed lines (wash). **A** Comparison of apoplastic roGFP2-Orp1 re-oxidation rates in *P. patens* and *A. thaliana* after pre-reduction with 5 mM TCEP (blue inverted triangles), with untreated samples serving as physiological controls (phys., green circles), n = 5. **B** Comparison between inherent re-oxidation rate (blue inverted triangles) and re-oxidation by adding 10 mM H_2_O_2_ after washing (red triangle), n = 4 for *P. patens*, n = 5 for *A. thaliana*.

As re-oxidation rates of roGFP2-Orp1 were lower than expected *in vivo*, we investigated if activation of apoplastic O_2_^·-^ production via RBOHs would increase sensor oxidation rates. To this end, we used *A. thaliana* leaf discs and the peptide elicitor *flg22* (**Fig. 4**) (Felix *et al*., 1999). First, we confirmed that the *flg22*-triggerred immune response is still functional after pre-reducing Col-*0* leaf discs using TCEP by luminol assays (**Fig. 4A**). Assessing roGFP-Orp1 fluorescence under the same conditions, we noticed that addition of *flg22* leads to an increase in autofluorescence after excitation of fluorescence at 400 nm that was detectable in control leaf discs from WT without sensor expression (**Fig. 4B** grey overlay). This massive increase in autofluorescence interfered with fluorescence-based ratiometric roGFP2 read-out c. 5 h after addition of *flg22*. We thus restricted our further analysis of the roGFP2-Orp1 400/480 fluorescence ratio to the first 4 h after addition of *flg22*, that did not show an increase of autofluorescence in WT (**Fig. 4C&D**).

**Figure 4:**
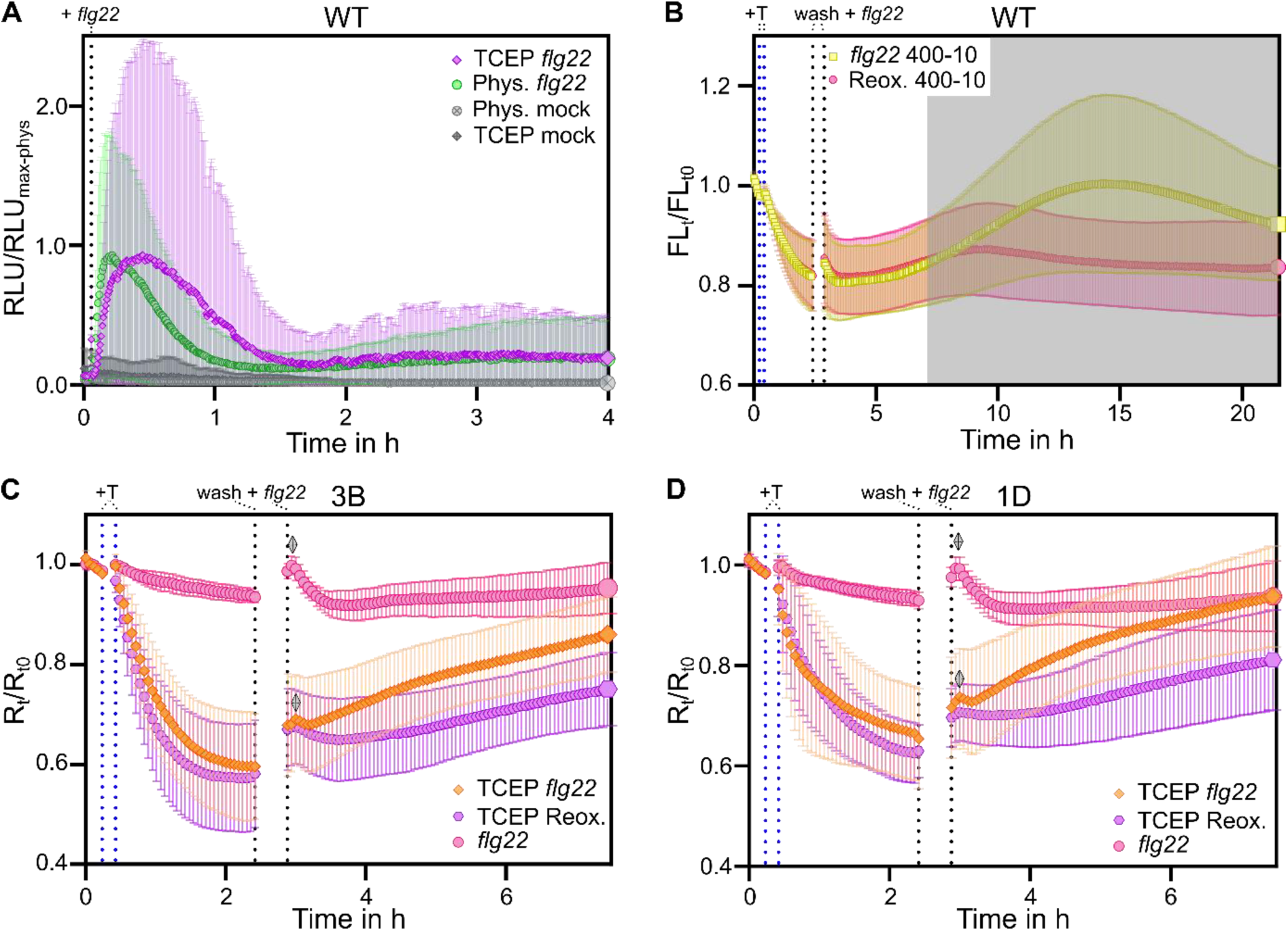
ROS burst after reduction of apoplastic roGFP2-Orp1. The luminescence (A) and fluorescence (B, C, D) of *A. thaliana* leaf discs expressing CHI_sp_-roGFP2-Orp1 (C, D) and Col-*0* (A, B) leaf discs were measured in a plate reader. **B, C, D**: The blue dashed lines indicate the addition of 5 mM TCEP. The black dashed lines indicate the wash out of TCEP and the addition of 1 µM flg22. **A** Normalised RLU of a luminescence assay plotted against the time. Samples were treated with 5 mM TCEP for 2 hours before the measurement (purple and grey diamonds), whereas control samples were kept in imaging buffer (green and grey circles). For the measurement, imaging buffer containing 20 µM L-012 and 8 µg/ml HRP1 was used. The dashed line indicates addition of 1 µM *flg22* or buffer, n = 29. **B** Autofluorescence profile in Col-*0* normalised over the initial intensity before treatment, plotted against the time. Each sample was treated with 5 mM TCEP whereas only part of the samples was treated with 1 µM *flg22* (yellow squares), n = 12. **C, D** Graphs depict the 400-10/482-16 ratio of apoplastic roGFP2-Orp1 in line 3B and 1D, plotted against the time. The ratio was normalised over the initial ratio before addition of treatment (R_t_/R_t0_). Depicted are samples first treated with TCEP and with *flg22* (orange diamonds), samples not treated with TCEP, but *flg22* (pink circles) and samples only treated with TCEP (purple hexagons). A transitory peak after addition of *flg22* is indicated by the two grey diamonds; n = 20.

In contrast to a control without addition of *flg22*, pre-reduced roGFP2-Orp1 re-oxidised during the first 3h after *flg22* addition, suggesting an increased H_2_O_2_-based oxidation rate in the apoplast. A second control sample, containing leaf discs expressing secreted roGFP2-Orp1 that were not pre-reduced showed a slight oxidative peak after addition of *flg22*, as well as the pre-reduced sample (grey diamonds in **Fig. 4C&D**). This transitory peak might be caused by the manual addition of *flg22*, and opening/closing of the plate reader, or represent a rapid transitory oxidation event, that is also affecting roGFP-Orp1 without pre-reduction. In summary, the elicitor-triggered immune response led to more rapid sensor re-oxidation in the apoplast of *A. thaliana* leaf discs, which was measurable via fluorescence read-out of roGFP2 in the first hours after initiation of an oxidative burst. As extracellular ROS are also important in a developmental context, we next investigated roGFP2-Orp1 signal both from an intracellular as well as an extracellular perspective in tip-growing gametophytic tissues. To this end, we compared both cytosolic and secreted roGFP2-Orp1 in *P. patens* protonema and *N. tabacum* pollen tubes (**Fig. 5, Figs. S12, S13**).

**Figure 5:**
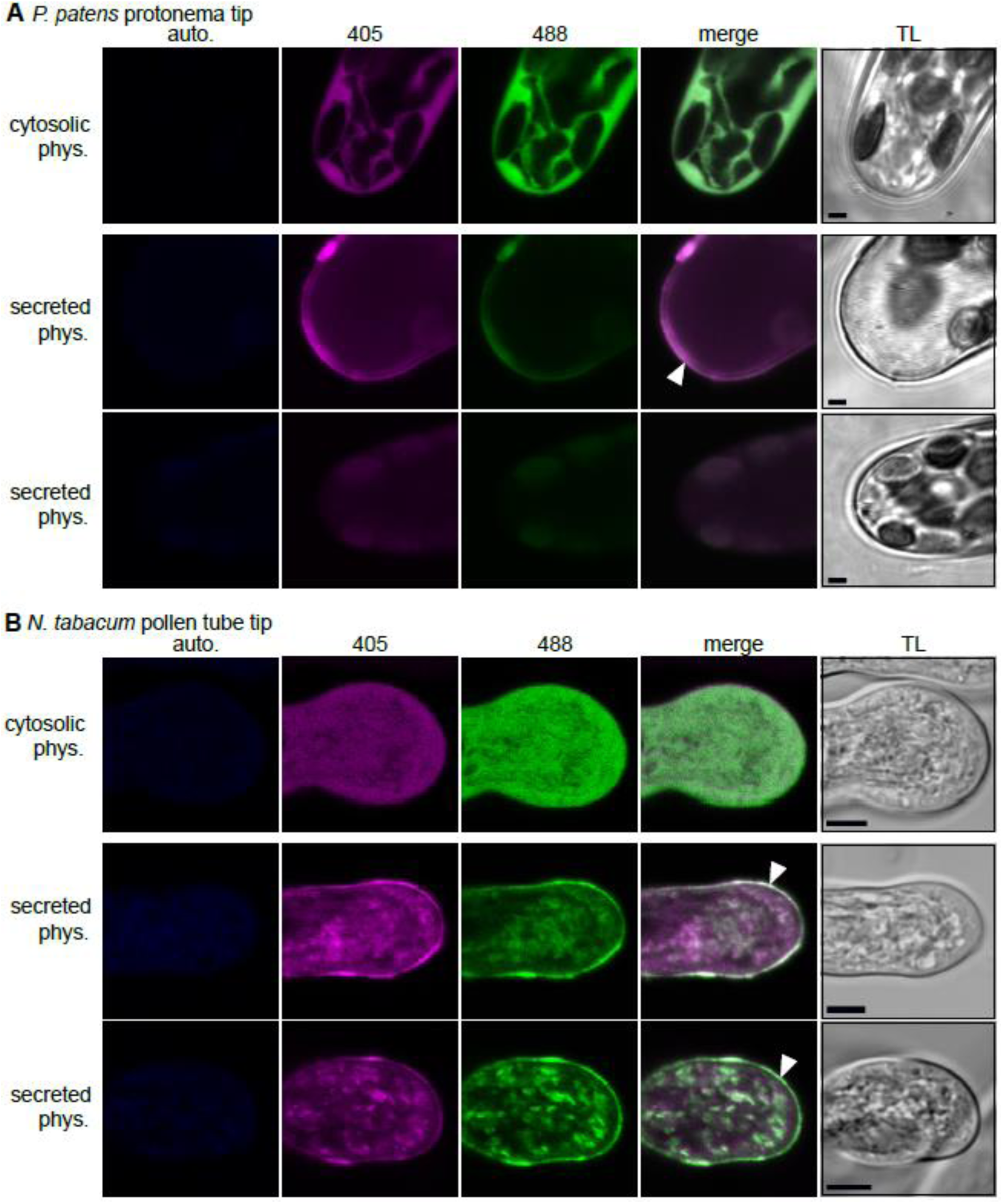
Comparison of cytosolic vs. secreted roGFP2-Orp1 during tip growth in *P. patens* protonema and *N. tabacum* pollen tubes. **A**, **B** Exemplary confocal images showing tips *P. patens* protonema tip cells and *N. tabacum* pollen tubes expressing cytosolic (upper panels) or secreted (middle and lower panels) *roGFP2-Orp1* under physiological (phys.) conditions. Most protonema tip cells show no tip roGFP2-Orp1 signal (A, lower panel) with few exceptions (A, middle panel). Autofluorescence (auto.) elicited after excitation at 405 nm is depicted in blue (λem: 425 - 475 nm), roGFP2 signal after excitation at 405 nm in magenta (λem: 509 - 535 nm) and the roGFP2 signal after excitation at 488 nm in green (λem: 509 - 535 nm). The 405/488 fluorescence ratio is shown as merge of both roGFP signals (merge); TL, transmitted light; arrowhead point to extracellular signals; scale bars = 2 µm (A), 5 µm (B).

We found that signal intensity and distribution of the secreted roGFP2-Orp1 differs between the species. In the apoplast of *P. patens* protonema tissue, most tip-growing meristematic cells showed no roGFP2-Orp1 accumulation in the apoplast, with few exceptions (**Fig. 5A, Fig. S12A**). In contrast, in the transition from tip growth to emerging gametophore buds, the apoplastic sensor signal was more intense and homogenous (**Fig. S12B**). Assessing transiently transformed *N. tabacum* pollen tubes, roGFP2-Orp1 signal was visible in secretory vesicles as well as in the apoplastic region (**Fig. 5B, Fig. S13**). Sensor calibration of *N. tabacum* pollen tubes revealed that secreted roGFP2-Orp1 showed a trend towards a lower, but not significantly different, 405/488 ratio compared to a completely oxidised control (**Fig. S13C**). Investigating cytosolic roGFP2-Orp1, we found an evenly reduced cytosolic signal in both protonema and pollen tubes (**Fig 5**). Comparing cytosolic and secreted roGFP2-Orp1 signals using the same microscopy settings for each cell type, secreted roGFP-Orp1 in protonema showed a clear oxidative shift compared to the cytosolic signal (**Fig. 5A, Fig. S12**). However, pollen tube tips were lacking this contrast and instead showed potentially partially reduced roGFP2-Orp1 (**Fig. 5B**).

## DISCUSSION

### Observed heterogeneity of roGFP2-Orp1 levels and redox states

In this study we generated and characterised stable transgenic plant lines secreting the redox sensor roGFP2-Orp1, using the model bryophyte *P. patens* and the model flowering plant *A. thaliana*. Bryophytes are the sister clade to vascular plants, undergoing separate evolution since c. 480 Mio. years (Lang *et al*., 2010; Donoghue *et al*., 2021). Mosses display a dominant gametophytic generation, with juvenile tip-growing filamentous protonema and leafy gametophores (Kofuji & Hasebe, 2014; Müller *et al*., 2016; Falz & Müller-Schüssele, 2019). By using species-specific signal peptides, we were able to mediate complete secretion of roGFP2-Orp1 in both model species, enabling further specific studies of sensor levels and redox steady state in the apoplast. We found that sensor levels differed between tissues for both species. In *P. patens*, protonema displayed lower roGFP2-Orp1 levels, with the highest fluorescence at perpendicular walls and only few tip cells showing apoplastic roGFP2-Orp1 signal at cell tips. The used endogenous Pp*Actin5* promoter drives constitutive expression, as evidenced by the stable transgenic lines expressing cytosolic roGFP2-Orp1. Thus, the overall low level of sensor signal in protonema cells is likely due to diffusion of secreted roGFP2-Orp1 into the surrounding media through protonema cell walls. This property of protonema is already known and used for biotechnological applications to secrete protein biopharmaceuticals into the surrounding media for downstream processing (Reski *et al*., 2015). In contrast, roGFP2-Orp1 homogenously labelled the apoplastic space in gametophore buds and leaflets, that possess a cuticle (Renault *et al*., 2017). Here, either cell wall density or the cuticle might limit the diffusion of secreted soluble particles out of the tissue. The Pp*Actin5* promoter was also active in moss gametangia, showing partially heterogenous signal with zones of high secretion and/or low diffusion. In comparison to *P. patens*, we observed higher heterogeneity of roGFP2-Orp1 fluorescence levels in *A. thaliana* within the same tissue. We currently interpret the higher fluorescence signals in lobes of pavement cells as well as in the meristematic/transition zone in root tips as indications of high secretion rates and/or low apoplastic diffusion rates. Punctate signals in mesophyll correlated with contact points between cells in spongy mesophyll, also suggesting high secretion or low apoplastic protein diffusion for these zones.

Ratiometric analysis of roGFP2-Orp1 fluorescence in its physiological state revealed mostly oxidised sensor protein, as expected and previously tested for the extracellular environment (Arnaud *et al*., 2023). The thiol-switch on roGFP2 has a consensus midpoint potential of-280 mV (pH7), that allows for sensitive measurement of redox changes in +/-35 mV of this value (Müller-Schüssele *et al*., 2021b). Thus, roGFP2 should be fully oxidised in the extracellular space with an estimated *E*_GSH_ less negative than-240 mV. Yeast Orp1 contains a peroxidatic cysteine (pK_a_ of 5.1 (Ma *et al*., 2007)) that, via its thiolate state, attacks H_2_O_2_, leading to sulfenylation and subsequent disulfide formation. This disulfide is then transferred by a dithiol/disulfide exchange mechanism to the fused roGFP2 moiety (Gutscher *et al*., 2009). Apoplastic pH in plants can rapidly change and has been measured as 6.3 in *A. thaliana* using the secreted pHluorin fluorescent sensor protein, with other values in literature ranging from c. 5 to 7 and potential spatio-temporal changes of up to 2 pH units (Gao *et al*., 2004; Martinière *et al*., 2013). Using fluorescent protein-based ‘Acidin’ biosensors, freely diffusing apoplastic sensor indicated a pH of c. 4.5 and showed very similar localisation pattern in lobes of pavement cells, compared to secreted roGFP2-Orp1 in this study (Moreau *et al*., 2022). In low pH (below c. 5) fluorescence after excitation at 488 nm (B-band) will be quenched in GFP variants (containing S65T) (Elsliger *et al*., 1999; Hanson *et al*., 2004). In contrast, absorption after excitation at 405 nm (A-band) is increased in lower pH (Elsliger *et al*., 1999), which will result in fluorescence in the case of roGFP2 (Hanson *et al*., 2004). In consequence, roGFP2 oxidation state can be measured in the pH range expected, in comparison to the respective *in vivo* calibration. We were able to calibrate apoplastic roGFP2-Orp1 using thiol-specific reductants. The dynamic measurement range (δ) in our hands was smaller than the δ6.1 reached with purified sensor or δ6.5 of cytosol and δ4.1 mitochondria-targeted sensor *in vivo* (Nietzel *et al*., 2019). It is already known that roGFP2-Orp1 δ decreases in lower pH (Nietzel *et al*., 2019). Lower δ can additionally be caused by low signal-to-noise ratios, non-optimal settings, difficulties in administering reduction and oxidation agents for *in vivo* calibration and/or interfering autofluorescence in different tissues as well as full or partial cleavage by local proteases.

Assessing physiological redox states of secreted roGFP2-Orp1 by both confocal microscopy as well as plate reader-based fluorescence read-out suggested that a minor fraction of sensor molecules might exist in reduced state, especially in zones of high sensor signals. This finding needs to be interpreted cautiously, as there are several possible explanations for our observations. Expressing a thiol-containing sensor such as roGFP2-Orp1 under a high-level constitutive promoter might disturb ER redox homeostasis, causing reductive stress (Ugalde *et al*., 2022a), resulting in secretion of proteins with reduced thiols. Additionally, irreversible post-translational modification of the redox-sensitive sensor cysteines might create non-responsive pseudo-reduced sensor proteins. However, as we were able to conduct sensor calibration within the described dynamic range, our results point to spatio-temporal zonation in the apoplast, creating areas where the redox steady state of protein cysteinyl groups (with similar characteristics to the protein sensor) is not completely oxidised. Mechanistically, possible reduction mechanisms affecting thiols in the apoplast are unclear to date, while both H_2_O_2_ and/or GSSG would mediate disulfide formation with slow rates non-enzymatically (Deponte, 2017).

Interestingly, we observed differences when comparing intracellular to secreted roGFP2-Orp1 signals in *P. patens* protonema to pollen tubes of *N. tabacum* that may relate to a different secretion/diffusion balance. Both cell types grow by tip growth (Kost *et al*., 1999; Menand *et al*., 2007; Bibeau *et al*., 2021), although the in vivo growth speed differs by several orders of magnitude, c. 4-10 µm h^-1^ in chloronema (Menand *et al*., 2007) vs. c. 300 µm h^-1^ in pollen tubes of *N. tabacum* (Geitmann *et al*., 1996). Although the *N. tabacum* growth rate is considered slow compared to the 10-20,000 µm h^-1^ of other angiosperms (Williams, 2012). The main role of short-lived pollen tubes is synthesis and assembly of cell wall components (Chebli *et al*., 2012) to extent themselves, bringing the sperm cells to ovules, powered by massive secretion at the apex. Notably, both investigated cell types do not possess cuticles but show different retention of secreted roGFP2-Orp1 around themselves. It is unclear if different cell wall properties could mediate this effect. At the apex, pollen tubes form a soft single layer cell wall with esterified pectin, cellulose and callose appearing with a distance of 5 – 30 µm behind the apex (Ferguson *et al*., 1998; Chebli *et al*., 2012; Wang *et al*., 2013). The pollen tube cell wall has evolved to be highly flexible, yet resistant to environmental stress, while balancing turgor pressure and fast directional growth (Chebli *et al*., 2012; Vogler *et al*., 2013; Wang *et al*., 2013). To a certain extent, the ability to store secreted proteins might contribute to sporophyte/gametophyte and gametophyte/gametophyte interactions as e.g. dissolving cuticle on female organs, growing through the extracellular matrix of transmitting tracts and signalling at pollen tube reception in ovules (Ingram & Nawrath, 2017; Becker *et al*., 2025). The redox state analysis of apoplastic roGFP2-Orp1 needs to be interpreted with caution, as ratiometric analysis of *N. tabacum* pollen tubes is challenging due to the fast growth and has been performed in transiently transfected cells growing in vitro. Showing the direct difference between cytosolic and secreted roGFP2-Orp1 in both protonema and pollen tube tips (**Fig.5**), we consistently observe no redox-gradients sensed by neither the cytosolic nor the secreted version of roGFP2-Orp1. This is contrasting to a previous data set using the pH-sensitive variant of the H_2_O_2_ sensor HyPer that detected a potential increase in sensor oxidation in the shank compared to the tip region (Boisson-Dernier *et al*., 2013), which might be explained as a result of a pH-gradient instead. Incomplete sensor oxidation in the apoplast of *N. tabacum* pollen tube apices may be caused by the massive secretory load and raises interesting questions regarding the local redox environment around pollen tubes.

### What can we learn from secreted roGFP2-Orp1 regarding apoplastic redox dynamics?

To investigate oxidation rates for protein cysteinyl residues *in vivo*, we pre-reduced samples and monitored sensor re-oxidation rates. Notably, after reduction with either a cell-permeable or a non-cell-permeable reducing agent for thiols, roGFP2-Orp1 redox state was stable for c. 20 hours of monitoring in *P. patens* gametophores, and only slowly re-oxidised in the time frame of hours in leaf discs of *A. thaliana* (**Fig. 3**, **Fig. S9**). In contrast, re-oxidation after exogenous addition of H_2_O_2_ was fast, demonstrating responsiveness of roGFP2-Orp1. Without the presence of enzymatic catalysis via glutaredoxins (GRX), equilibration of proteins cysteinyl residue redox steady state and local *E*_GSH_ is slow (Gutscher *et al*., 2008; Deponte, 2017; Bohle *et al*., 2024b). Thus, *E*_GSH_-dependent roGFP2 oxidation rates would potentially fit the observed sensor behaviour in the apoplast. In contrast, H_2_O_2_-mediated oxidation of Orp1 is kinetically fast, due to higher rate constants, even in the pH range expected for the apoplast. Thus, slow sensor re-oxidation indicates low H_2_O_2_-mediated oxidation rates in the apoplast under physiological conditions. Exact local H_2_O_2_ levels cannot be directly derived from sensor read-out, but roGFP2-Orp1 has been shown to sense as low as 0.1 µM H_2_O_2_ *in vitro* (Gutscher *et al*., 2009; Nietzel *et al*., 2019). According to current knowledge, RBOH activity causes a rapid extracellular increase of H_2_O_2_ levels. RBOHs are transmembrane proteins that can transfer electrons from cytosolic NADPH to extracellular oxygen, generating O_2_^·-^ anions and in consequence H_2_O_2_ and O_2_ by (spontaneous) dismutation. Extracellular H_2_O_2_ is implicated in multiple processes, including signalling and pathogen defence. As it is known that treatment with the bacterial elicitor *flg22* in *A. thaliana* leads to RBOHD-dependent apoplastic oxidative bursts within seconds to minutes (Miller *et al*., 2009; Nietzel *et al*., 2019), we investigated the effect of *flg22*-treatment on secreted roGFP2-Orp1 re-oxidation rates. As autofluorescence can rapidly increase in consequence to plant immune responses (Bohle *et al*., 2024a), roGFP2 fluorescence-based read-out, especially after excitation in the UV range (e.g. 405 nm), needs to be carefully interpreted. Thus, the increase in autofluorescence bleed-through into the 400ex./520em. roGFP2 channel that we observed prohibits conclusions regarding the redox state after more than c. 4 hours after *flg22* treatment. However, we found increasing oxidation rates for pre-reduced roGFP2-Orp1 in the first hours after triggering RBOH activity via *flg22*. Compared to exogenous addition of 10 mM H_2_O_2_, re-oxidation rates after triggering an oxidative burst were slower. Using secreted roGFP-Orp1, we observed a minor transitory oxidation peak after *flg22* addition, which may be an artefact due to the manual addition of *flg22*. We observed a time lag of c. 1 h until apoplastic roGFP-Orp1 started oxidising at a stable rate. Based on the known phases of ROS generation during an immune response (Ngou *et al*., 2021; Arnaud *et al*., 2023), an immediate increase in apoplastic H_2_O_2_ levels was expected. This raises the question why protein-based H_2_O_2_ sensing in the apoplast differs from luminol-based assays. Importantly, also intracellular redox biosensing revealed a time lag between luminol-based oxidative burst detection and intracellular oxidative changes (Nietzel *et al*., 2019; Ugalde *et al*., 2022b; Arnaud *et al*., 2023). Intracellular oxidation of cytosolic roGFP-Orp1 starts in the range of minutes to an hour after triggering an oxidative burst with *flg22* in *A. thaliana*, while H_2_O_2_-detection with luminol-based assays indicates a rapid peak in the first 20 min (Nietzel *et al*., 2019; Arnaud *et al*., 2023). Arnaud *et al*. (2023) revealed that the bi-phasic cytosolic roGFP2-Orp1 oxidation observed in response to *flg22*-triggered immune signalling was largely unchanged in *rbohd* plants. Thus, while luminescence assays clearly evidence RBOHD activity, RBOHD activity is not necessary for intracellular oxidation dynamics in response to *flg22*. In this context, our results indicate that extracellular oxidative changes occurring downstream of RBOH activity are not as massive as suggested by luminol-based assays, or more specific.

In conclusion, we found lower than expected oxidation rates for extracellular protein cysteinyl residues in *A. thaliana* that are even lower to absent in the model bryophyte *P. patens*. Our work provides first indications to the timeline of increasing apoplastic H_2_O_2_ levels in consequence to activating an immune response, as directly measured with a protein-based redox sensor. However, it is unclear to date if cysteinyl oxidation rates in the extracellular space are biologically relevant. In animal cells, dynamic redox steady states in the extracellular space are linked to several processes such as proliferation, differentiation and cell death (Banerjee, 2012). In land plants, the RBOH gene family has expanded and diversified, playing important roles in immunity, but also during reproduction, e.g. in pollen tip growth (Kaya *et al*., 2014; Mhamdi & Van Breusegem, 2018). How RBOHs fulfil their specific and diverse roles in creating local and dynamic apoplastic redox environments requires further investigations. In absence of interfering autofluorescence, roGFP2-based sensors may be used to investigate oxidation dynamics. This approach could be particularly interesting to resolve spatial differences in a tissue context. Detection of apoplastic redox dynamics by cysteine-based sensors remains challenging, due to the need of pre-reduction. In addition, secreted roGFP2-Orp1 (or other pH-stable fluorescent proteins) can provide valuable information regarding zones of secreted protein retention and diffusion barriers in different plant species and tissues.

## Acknowledgements

The authors are grateful for the funding from the DFG (Research Unit FOR 5098 ‘Innovation and Coevolution in Plant Sexual Reproduction’ (JI, PLZ, ST, SS, TD, SJMS)). SJMS is grateful for funding obtained from BioComp 3.0 ‘Dynamic Membrane Processes in Biological Systems’. We thank Aurora Martin (Molecular Botany, RPTU), Alexa Brox and Maria Homagk (Chemical Signalling, INRES, University of Bonn) for technical assistance. We are grateful to Anna-Lena Falz, Cloé Gadoud, Emma Müller, Sadia Tamanna, Ralf Pennther-Hager and Dominik Stehr for experimental support. We thank Markus Schwarzländer for providing plasmids and plant lines for cytosolic roGFP2-Orp1 expression, as well as useful discussion.

## Author Contributions

JI, OT and SJMS designed the research. JI, LZ, PLZ and ST performed experiments and analysed data. SJMS, OT, AJM, TD and SS supervised the research and provided resources. JI and SJMS wrote the manuscript with contributions from all authors. All authors approved the manuscript before submission.

## Data availability

Data acquired in this study are available in the Supporting Information of this article (Figs S1– S13). Requests for resources can be made to the corresponding author.

## Supporting Information

The following supplementary information is available:

*Fig. S1*. Transient expression of *CHI_sp_-roGFP2-Orp1* in pavement cells of *Nicotiana benthamiana* leaves.

*Fig. S2.* Creation of stable transgenic lines expressing secreted roGFP2-Orp1 in *P. patens* and *A. thaliana* (additional independent lines)

*Fig. S3.* Fluorescence signal distribution of apoplastic roGFP2-Orp1 in *P. patens* and *A. thaliana*

*Fig. S4.* Confocal images of AP1_sp_-roGFP2-Orp1 calibration in *P. patens*

*Fig. S5.* Confocal images of Ap1_sp_-roGFP2-Orp1 calibration in *P. patens* gametangia (line #131)

*Fig. S6.* Confocal images Ap1_sp_-roGFP2-Orp1 calibration in *P. patens* gametangia (line #19)

*Fig. S7.* Confocal images CHI_sp_-roGFP2-Orp1 calibration in *A. thaliana* leaf discs (line 3B)

*Fig. S8.* Confocal images of CHI_sp_-roGFP2-Orp1 calibration in *A. thaliana* leaf discs (line 1D)

*Fig. S9*. Re-oxidation rate of apoplastic roGFP2-Orp1 pre-reduced using DTT

*Fig. S10.* Re-oxidation rate of apoplastic roGFP2-Orp1 in *P. patens* and *A. thaliana* using varying doses of DTT

*Fig. S11.* Treatment with TCEP does not influence cytosolic roGFP2-Orp1 redox state in *A. thaliana*

*Fig. S12: Apoplastic roGFP2-Orp1 in P. patens protonema tip cells and young buds*

*Fig. S13 Comparison of roGFP2-Orp1 in the cytosol and in the apoplast of N. tabacum pollen tubes*.

