## Supplemental Figures1-13 for "The secreted redox sensor roGFP2-Orp1 reveals oxidative dynamics in the plant apoplast"

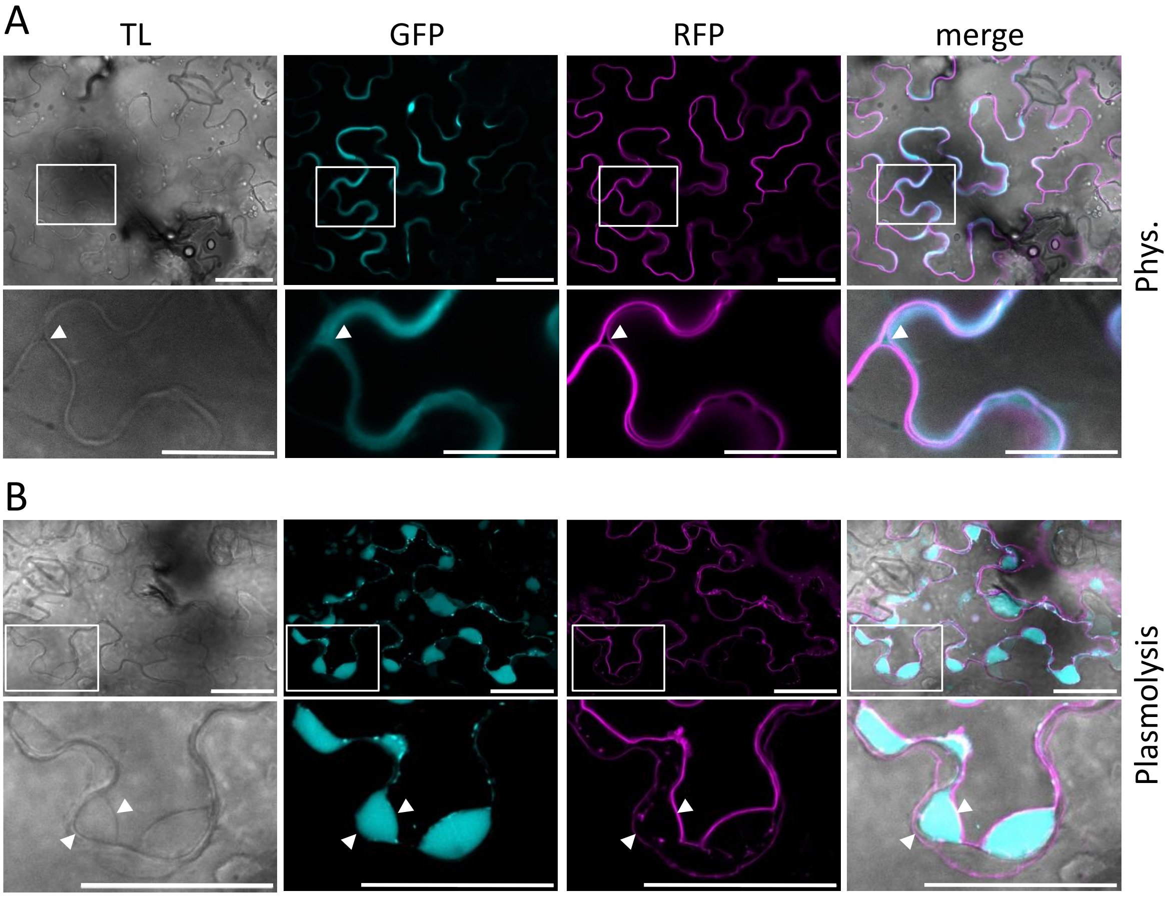


**Figure S1: Transient expression of *35S$_{pro}$-CHI$_{sp}$-roGFP2*-Orp1 in pavement cells of *Nicotiana benthamiana* leaves.**
To confirm secretion of roGFP2-Orp1, the sensor construct *35S$_{pro}$-CHI$_{sp}$-roGFP2-Orp1* was transiently co-expressed with the plasma membrane marker construct *35S$_{pro}$-TagRFP-T-RemA* in *N. benthamiana* leaf epidermal cells. Leaves were analyzed two days after infiltration with agrobacteria. Plasmolysis was induced by infiltrating 1 M sorbitol solution. From left to right: Transmitted light (TL), GFP (cyan), TagRFP-T (magenta) and merged channels. Lower panels show enlarged sections from the image above (boxed areas). Arrow heads in the lower panels point to the plasma membrane labelled by TagRFP-T-RemA. Scale bars =50 µm. **A** Cells expressing *CHI$_{sp}$-roGFP2-Orp1* (phys.). **B** Cells expressing *CHI$_{sp}$-roGFP2-Orp1* after plasmolysis; lower panel: close-up showing the separated of the plasma membrane detached from the cell wall and the localization of CHI$_{sp}$-roGFP2-Orp1 in the apoplast. Scale bars = 25 µm.

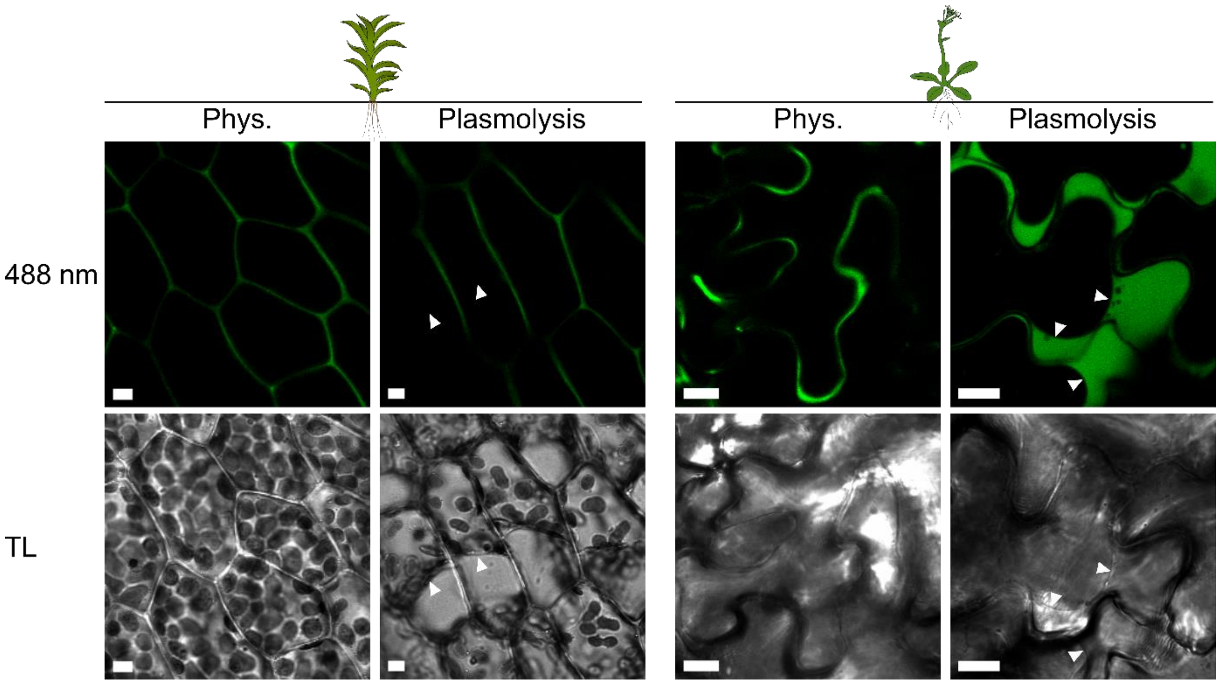


**Figure S2: Creation of stable transgenic lines expressing secreted roGFP2-Orp1 in *P. patens* and *A. thaliana* (additional independent lines)**
Exemplary confocal images of *P. patens* leaflets (line #19) and *A. thaliana* pavement cells (line 1D) stably expressing *AP1*$_{sp}$*-roGFP2-Orp1* or *CHI*$_{sp}$*-roGFP2-Orp1*, respectively. Plasmolysis was induced using 0.6 M Mannitol (*A. thaliana*) or 0.6 M NaCl (*P. patens*). Scale bars = 10 µm.

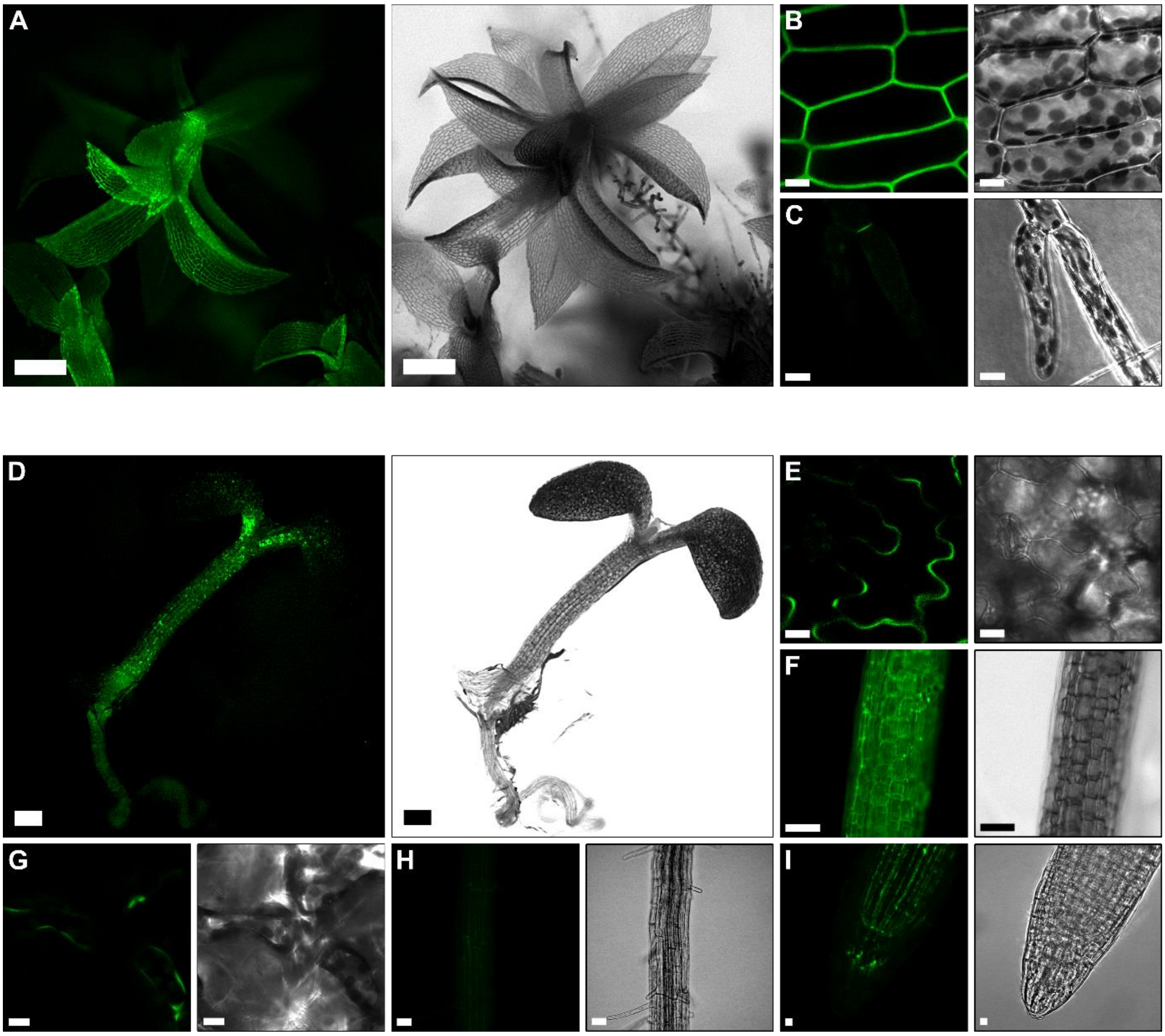


**Figure S3: Fluorescence signal distribution of apoplastic roGFP2-Orp1 in *P. patens* and *A. thaliana***

Overview of different *P. patens* tissues stably expressing *AP1$_{sp}$-roGFP2-Orp1* and *A. thaliana* tissues stably expressing *CHI$_{sp}$-roGFP2-Orp1*. Fluorescence signal of roGFP2 after excitation with blue light (480 or 488 nm) is depicted in green (left panels) and transmitted light or brightfield image in greyscale (right panels). **A** Overview of a gametophore, taken at a stereo microscope; scale bar = 100 µm. **B** shows cells of a leaflet. **C** shows protonema cells, B & C, were taken by CLSM; scale bar = 10 µm. **D** Overview of a seven-day-old *A. thaliana* seedling. **E** shows pavement cells of the cotyledons. **F** shows cells of the hypocotyl. **G** shows mesophyll cells of an adult leaf. **H** shows root with root hairs. **I** shows a root tip. D & F were taken at a stereo microscope; scale bars = 100 µm. E, G, H & I were taken using CLSM; scale bars = 10 µm.

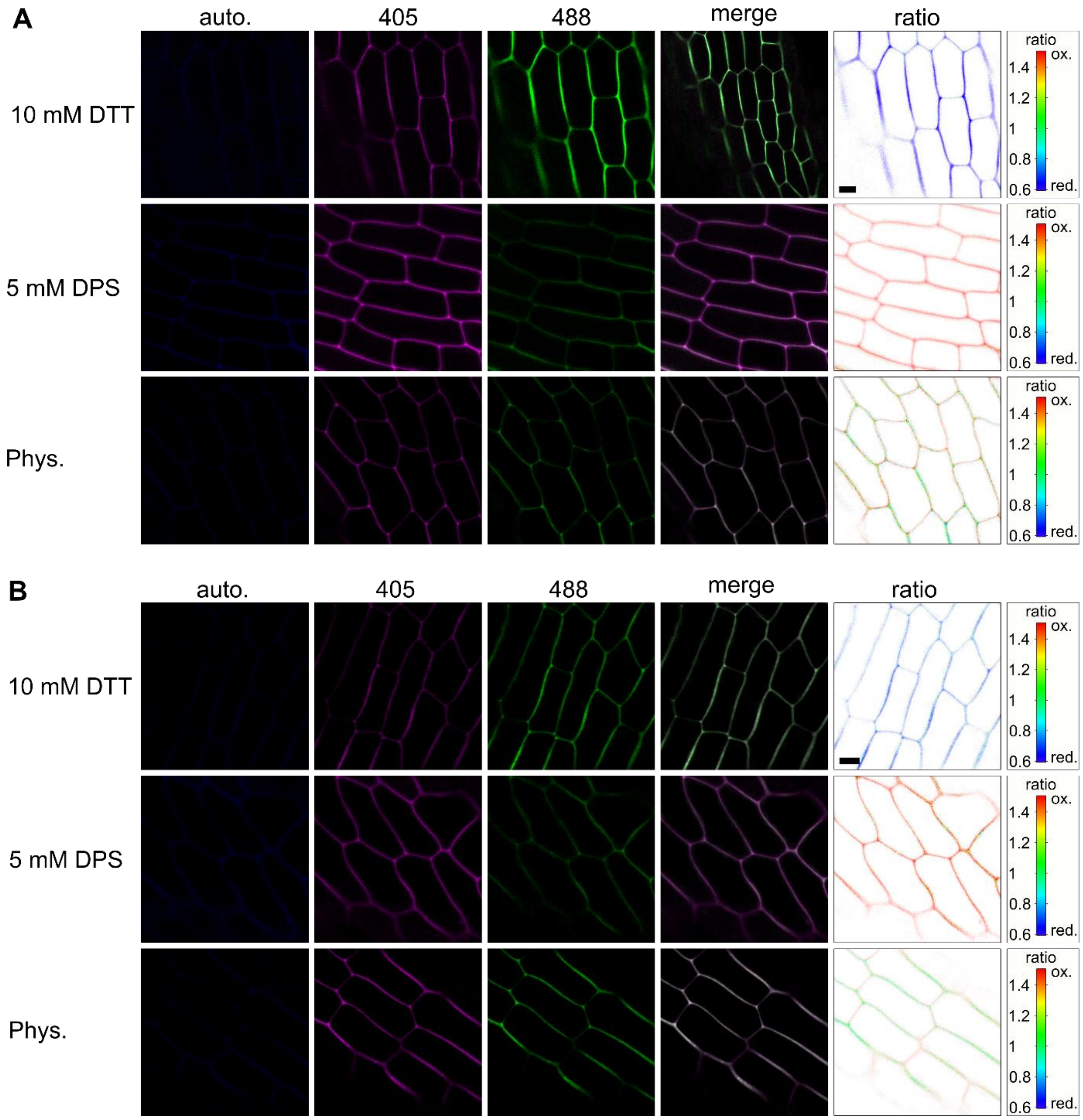


**Figure S4: Confocal images of AP1$_{sp}$-roGFP2-Orp1 calibration in *P. patens***
Sensor calibration is shown in two independent *P. patens* lines expressing *AP1$_{sp}$-roGFP2-Orp1*, #131 (A) and #19 (B). **A, B** Exemplary images showing individual fluorescence channels used as well as ratiometric analysis. Autofluorescence (auto.) elicited after excitation at 405 nm is depicted in blue (λem: 425 - 475 nm). The roGFP2 signal after excitation at 405 nm is depicted in magenta (λem: 509 - 535 nm) and the roGFP2 signal after excitation at 488 nm is depicted in green (λem: 509 - 535 nm). The 405/488 fluorescence ratio is shown as merge of both roGFP signals (merge) and as false-colour-coded ratio (ratio). DTT induces full sensor reduction whereas DPS induces full sensor oxidation. Phys.: physiological sensor oxidation state in untreated tissue. Scale bars = 10 µm.

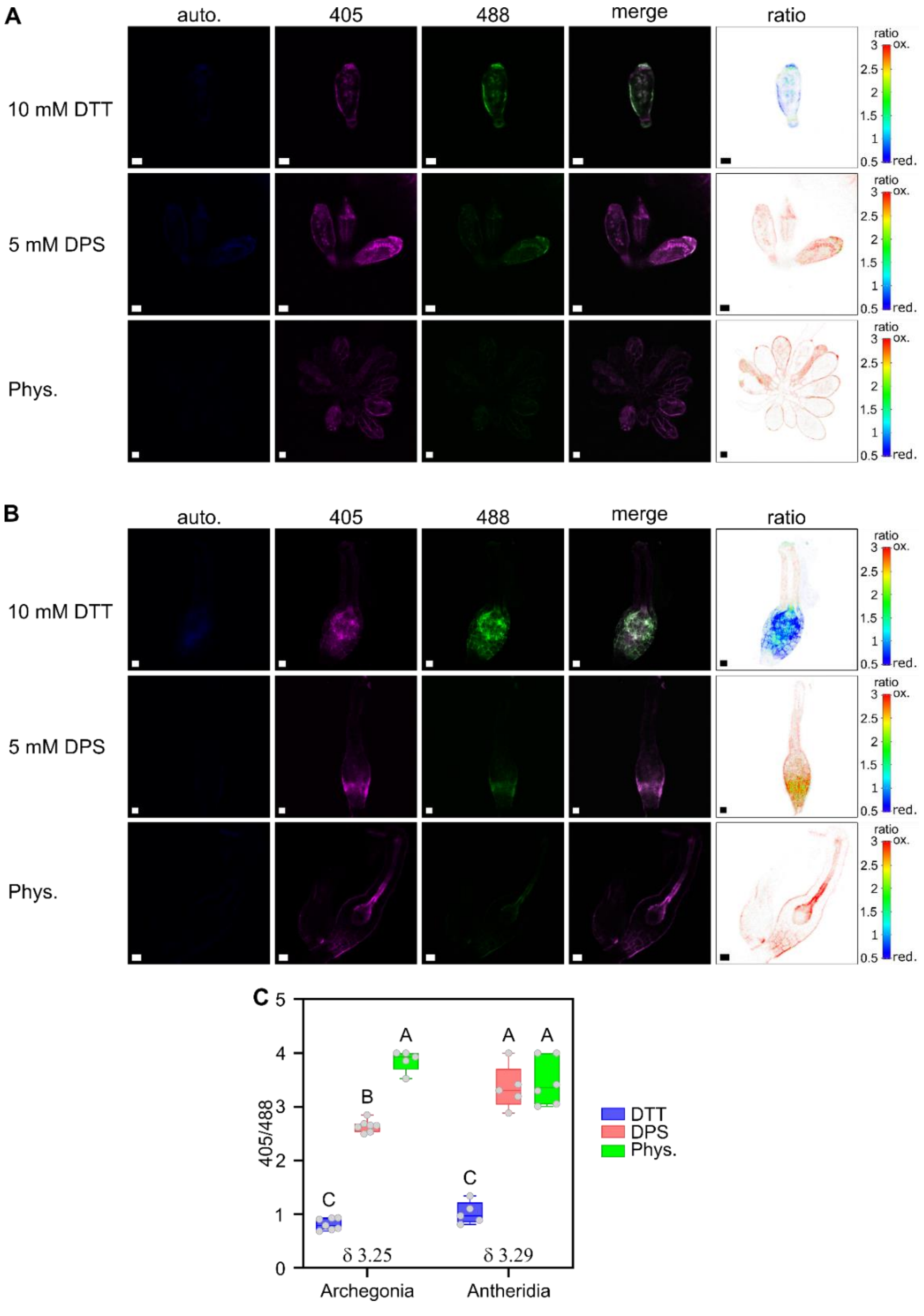


**Figure S5: Confocal images of Ap1$_{sp}$-roGFP2-Orp1 calibration in *P. patens* gametangia (line #131)**

Sensor calibration is shown for the line #131 in both gametangia: antheridia (A) and archegonia (B). **A**, **B** Exemplary images showing individual fluorescence channels used as well as ratiometric analysis. Autofluorescence (auto.) elicited after excitation at 405 nm is depicted in blue (λem: 425 - 475 nm). The roGFP2 signal after excitation at 405 nm is depicted in magenta (λem: 509 - 535 nm) and the roGFP2 signal after excitation at 488 nm is depicted in green (λem: 509 - 535 nm). The 405/488 fluorescence ratio is shown as merge of both roGFP signals (merge) and as false-colour-coded ratio (ratio). DTT induces full sensor reduction whereas DPS induces full sensor oxidation. Phys.: physiological sensor oxidation state in untreated tissue. Scale bar = 10 µm. **C** Box plots of redox ratio analysis using confocal microscopy images, measured dynamic range is indicated by δ. Boxes show 25th to 75th percentiles, line indicates the median, whiskers show min and max, individual data points are shown as circles, n = 5–7. Different letters indicate significant differences according to a two-way ANOVA with Tukey's multiple comparison post-hoc test, $p<0.05$.

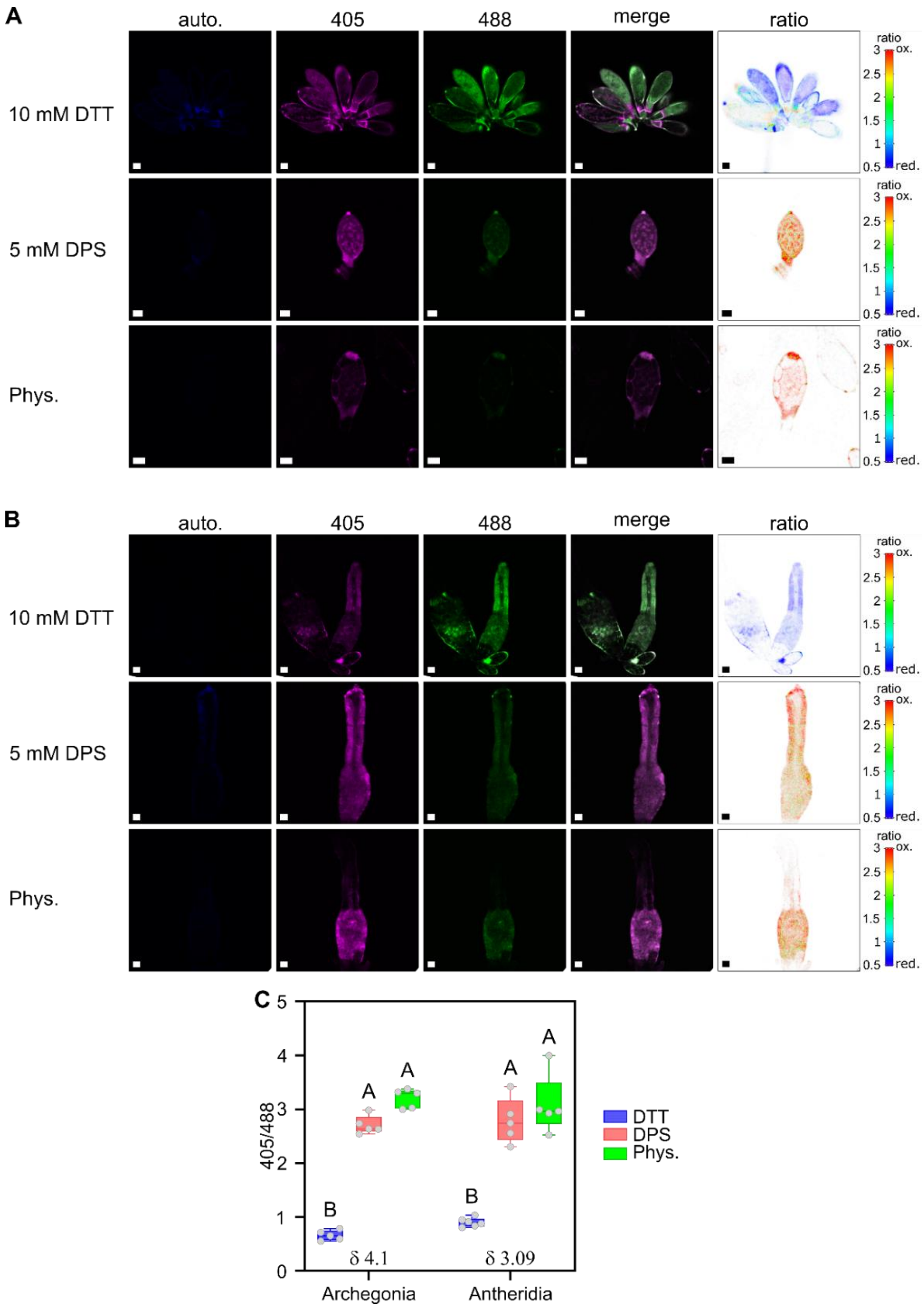


**Figure S6: Confocal images Ap1$_{sp}$-roGFP2-Orp1 calibration in *P. patens* gametangia (line #19)**

Sensor calibration is shown for the line #19 in both gametangia: antheridia (A) and archegonia (B). **A**, **B** Exemplary images showing individual fluorescence channels used as well as ratiometric analysis. Autofluorescence (auto.) elicited after excitation at 405 nm is depicted in blue (λem: 425 - 475 nm). The roGFP2 signal after excitation at 405 nm is depicted in magenta (λem: 509 - 535 nm) and the roGFP2 signal after excitation at 488 nm is depicted in green (λem: 509 - 535 nm). The 405/488 fluorescence ratio is shown as merge of both roGFP signals (merge) and as false-colour-coded ratio (ratio). DTT induces full sensor reduction whereas DPS induces full sensor oxidation. Phys.: physiological sensor oxidation state in untreated tissue. Scale bar = 10 µm. **C** Box plots of redox ratio analysis using confocal microscopy images, measured dynamic range is indicated by δ. Boxes show 25th to 75th percentiles, line indicates the median, whiskers show min and max, individual data points are shown as circles, n = 5-6. Different letters indicate significant differences according to a two-way ANOVA with Tukey's multiple comparison post-hoc test, $p<0.05$.

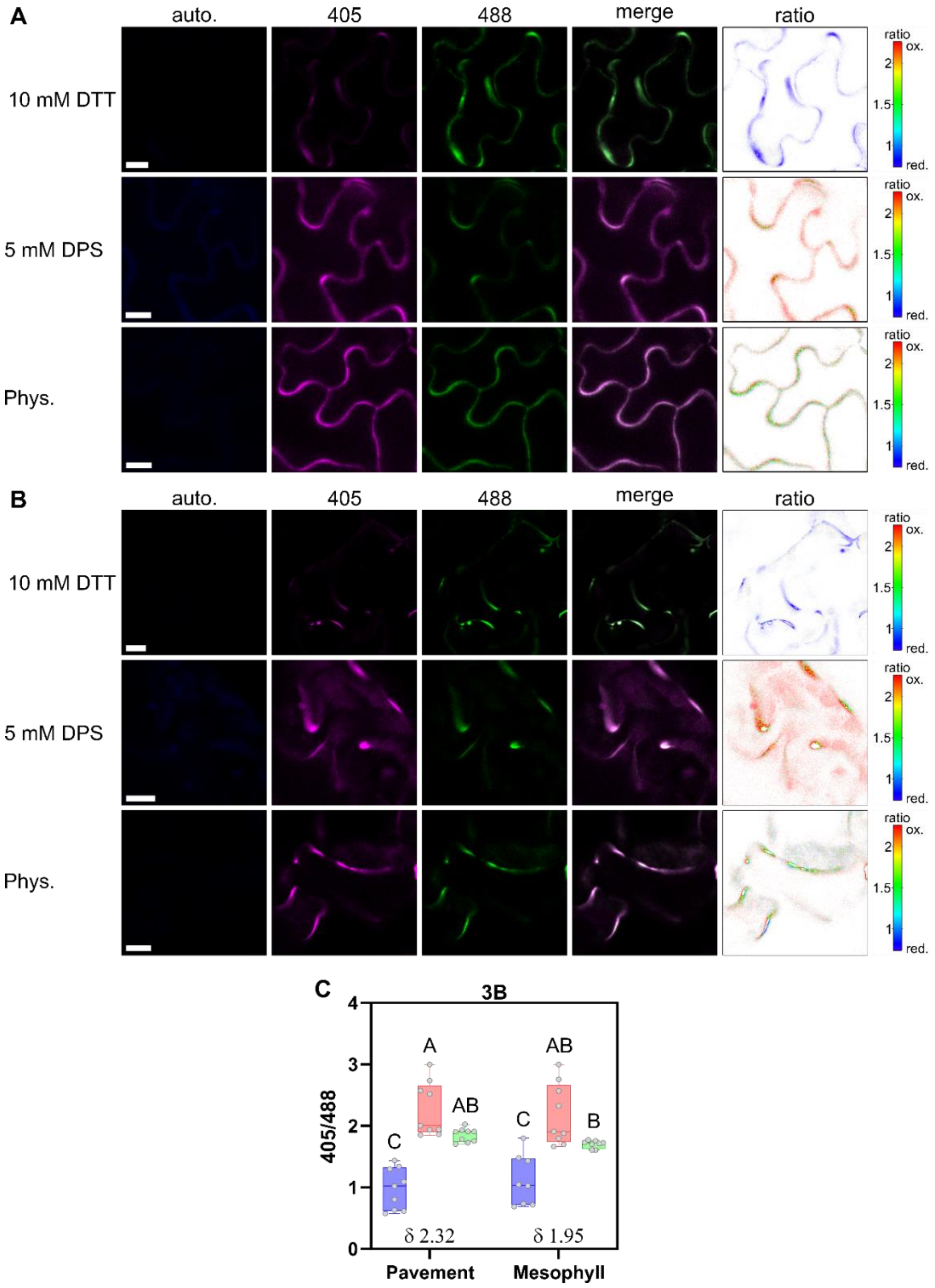


**Figure S7: Confocal images CHI$_{sp}$-roGFP2-Orp1 calibration in *A. thaliana* leaf discs (line 3B)**

Sensor calibration is shown for the line 3B in two different tissues, pavement cells (A) and mesophyll cells (B). **A, B** Exemplary images showing individual fluorescence channels used as well as ratiometric analysis. Autofluorescence (auto.) elicited after excitation at 405 nm is depicted in blue (λem: 425 - 475 nm). The roGFP2 signal after excitation at 405 nm is depicted in magenta (λem: 509 - 535 nm) and the roGFP2 signal after excitation at 488 nm is depicted in green (λem: 509 - 535 nm). The 405/488 fluorescence ratio is shown as merge of both roGFP signals (merge) and as false-colour-coded ratio (ratio). DTT induces full sensor reduction whereas DPS induces full sensor oxidation. Phys.: physiological sensor oxidation state in untreated tissue. Scale bar = 10 µm. **C** Box plots of redox ratio analysis using confocal microscopy images, measured dynamic range is indicated by δ. Boxes show 25th to 75th percentiles, line indicates the median, whiskers show min and max, individual data points are shown as circles, n = 8-9. Different letters indicate significant differences according to a two-way ANOVA with Tukey’s multiple comparison post-hoc test, $p<0.05$.

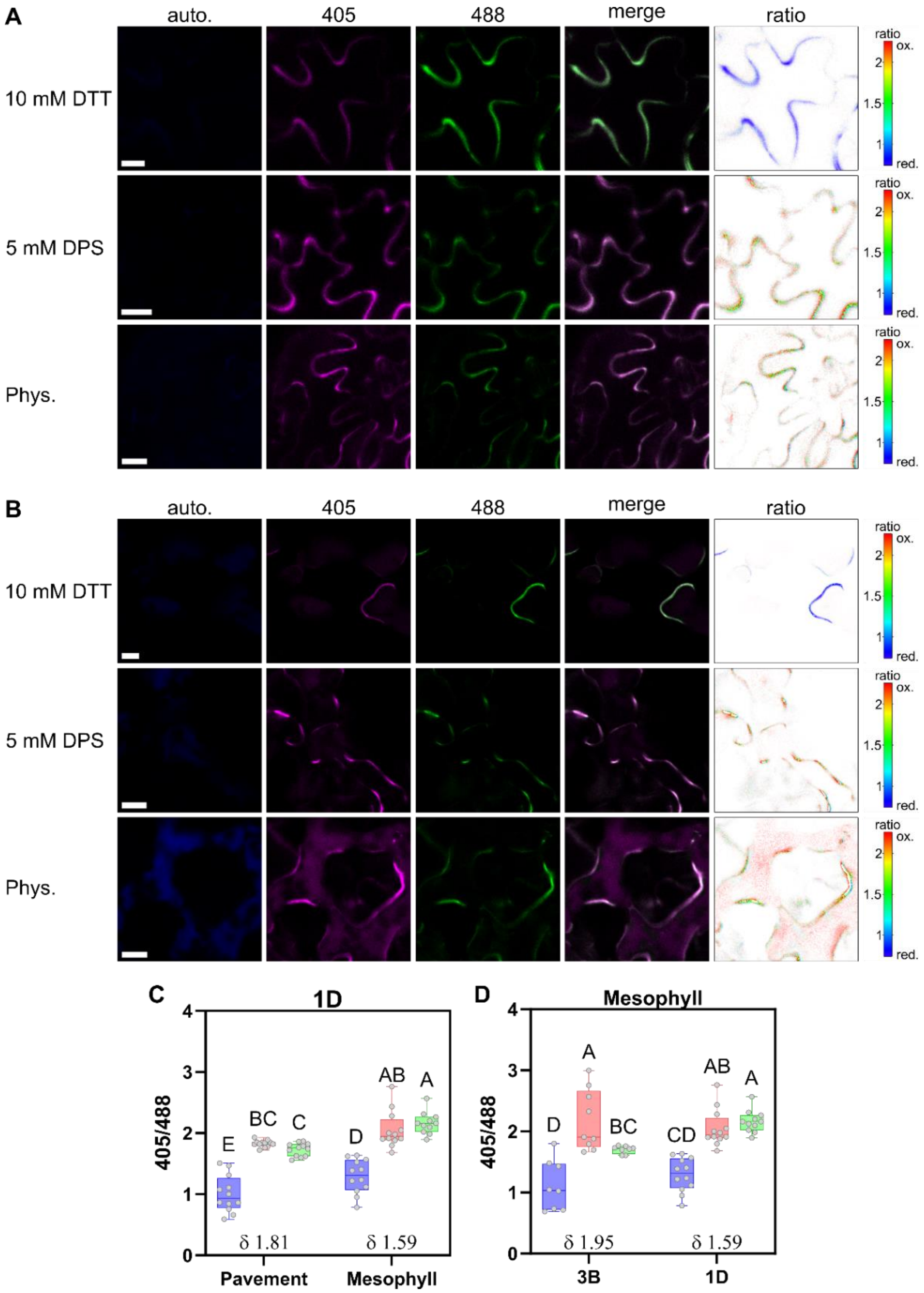


**Figure S8: Confocal images of $CHI_{sp}$-roGFP2-Orp1 calibration in *A. thaliana* leaf discs (line 1D)**

Sensor calibration is shown for the line 1D in two different tissues, pavement cells (A) and mesophyll cells (B). **A, B** Exemplary images showing individual fluorescence channels used as well as ratiometric analysis. Autofluorescence (auto.) elicited after excitation at 405 nm is depicted in blue (λem: 425 - 475 nm). The roGFP2 signal after excitation at 405 nm is depicted in magenta (λem: 509 - 535 nm) and the roGFP2 signal after excitation at 488 nm is depicted in green (λem: 509 - 535 nm). The 405/488 fluorescence ratio is shown as merge of both roGFP signals (merge) and as false-colour-coded ratio (ratio). DTT induces full sensor reduction whereas DPS induces full sensor oxidation. Phys.: physiological sensor oxidation state in untreated tissue. Scale bar = 10 µm. **C** Box plots of redox ratio analysis using confocal microscopy images, measured dynamic range is indicated by δ. Boxes show 25th to 75th percentiles, line indicates the median, whiskers show min and max, individual data points are shown as circles, n = 11-12. Different letters indicate significant differences according to a two-way ANOVA with Tukey's multiple comparison post-hoc test, $p<0.05$.

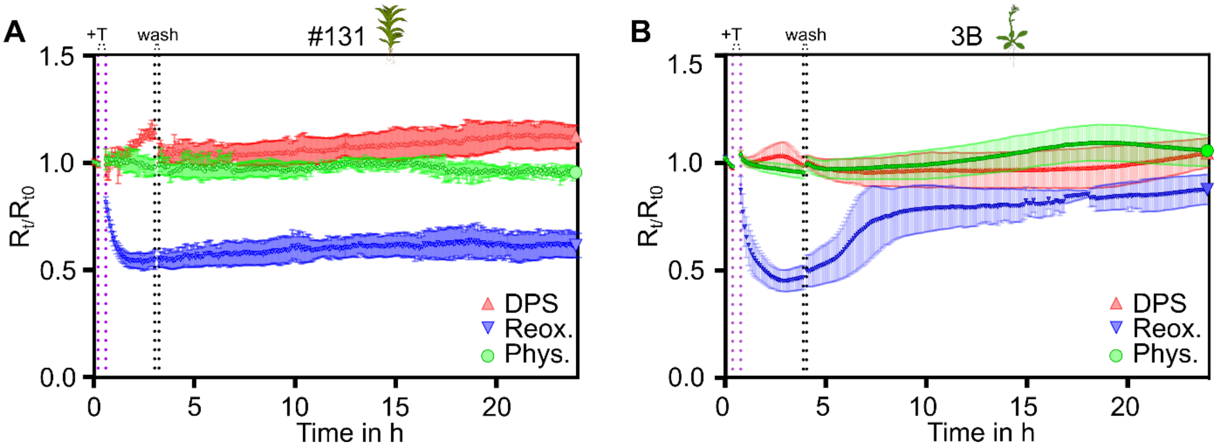


**Figure S9: Re-oxidation rate of apoplastic roGFP2-Orp1 pre-reduced using DTT**
Graphs depict 400-10/482-16 ratios, normalised to the ratio mean before treatment ($R_t/R_{t0}$), of apoplastic roGFP2-Orp1 measured as a time series using gametophores (*P. patens* line #131) or leaf discs (*A. thaliana* line 3B) in a multi-well plate reader. Start of treatment with 10 mM DTT is indicated by a purple dashed line (+T). After reaching plateau values the measurement was paused to remove and wash out the reducing agent, indicated by the two dashed lines (wash). **A** Apoplastic roGFP2-Orp1 re-oxidation rates in *P. patens* after pre-reduction with 10 mM DTT (blue inverted triangles), with untreated samples serving as physiological controls (phys., green circles) and samples treated with 5 mM DPS as oxidation controls (red triangles), n = 5. **B** Apoplastic roGFP2-Orp1 re-oxidation rates in *A. thaliana* after pre-reduction with 10 mM DTT (blue inverted triangles), with untreated samples serving as physiological controls (phys., green circles) and samples treated with 5 mM DPS as oxidation controls (red triangles), n = 5.

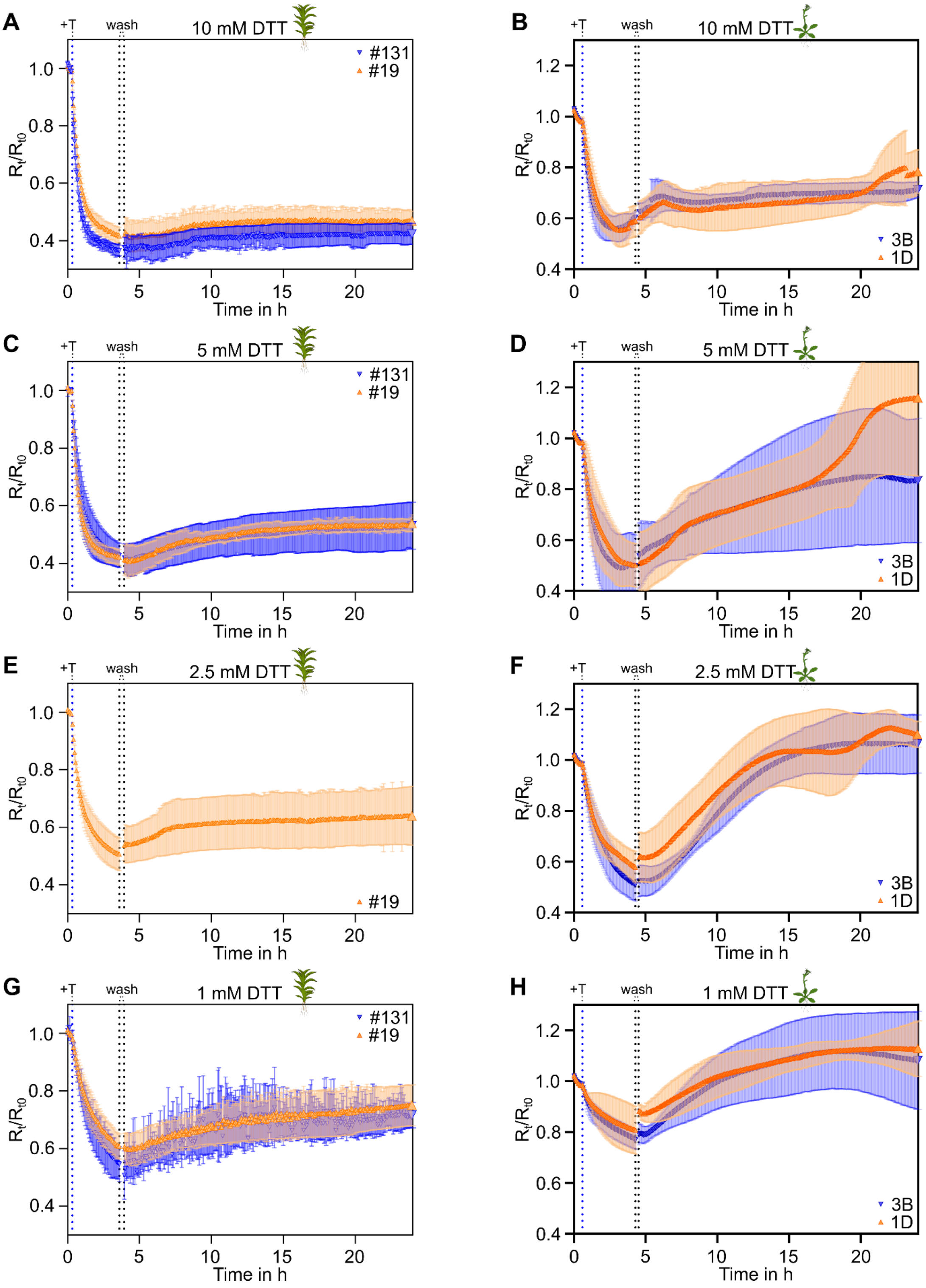


**Figure S10: Re-oxidation rate of apoplastic roGFP2-Orp1 in *P. patens* and *A. thaliana* using varying doses of DTT**

**A-H** Graphs depict 400-10/482-16 ratios, normalised to the ratio mean before treatment ($R_t/R_{t0}$) of apoplastic roGFP2-Orp1 measured as a time series using gametophores (*P. patens*) or leaf discs (*A. thaliana*) in a multi-well plate reader. Start of treatment with varying concentration of DTT is indicated by a blue dashed line (+T); DTT concentration used is depicted above panels. After maximal reduction was reached, the reductant was washed out, indicated by black dashed lines. A, C, G & E: *P. patens* gametophores of two independent lines # 131 (blue inverted triangle) and # 19 (orange triangle); n = 3-5. B, D, F, H: Leaf discs of two independent *A. thaliana* lines 3B (blue inverted triangle) & 1D (orange triangle); n = 5.

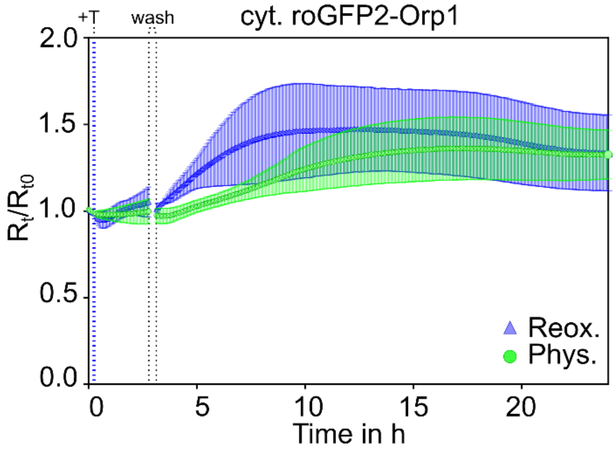


**Figure S11: Treatment with TCEP does not influence cytosolic roGFP2-Orp1 redox state in *A. thaliana***

Graphs depict 400-10/482-16 ratios, normalised to the ratio mean before treatment (Rt/Rt0). Fluorescence of leaf discs of a stable *A. thaliana* line expressing roGFP2-Orp1 in the cytosol (Nietzel et al., 2019) was measured, using a multi-well plate reader. Start of treatment with 5 mM TCEP is indicated by a blue dashed line (+T). After reaching plateau values the measurement was paused to remove and wash out the reducing agent, indicated by the two dashed lines (wash), n = 5. Data have been acquired in the same plate reader run depicted in Fig. 3A, right panel.

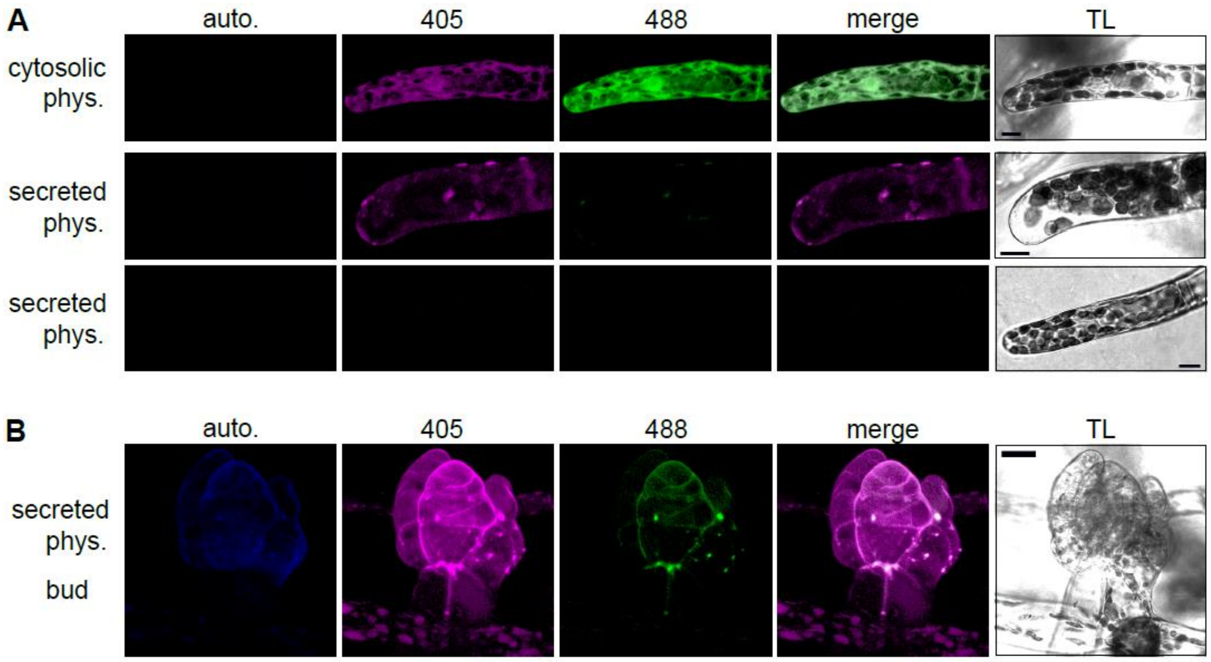


**Figure S12: Apoplastic roGFP2-Orp1 in *P. patens* protonema tip cells and young buds** Exemplary images showing individual fluorescence channels used as well as 405/488 merge and transmitted light (TL). Autofluorescence (auto.) elicited after excitation at 405 nm is depicted in blue (λem: 425 - 475 nm). The roGFP2 signal after excitation at 405 nm is depicted in magenta (λem: 509 - 535 nm) and the roGFP2 signal after excitation at 488 nm is depicted in green (λem: 509 - 535 nm). The 405/488 fluorescence ratio is shown as merge of both roGFP signals (merge); fluorescence images are maximum intensity projections from confocal z-stacks; phys.: physiological sensor oxidation state in untreated tissue. Scale bars = 10 µm (A), 20 µm (B). **A** Intracellular vs. extracellular perspective in tip cells of protonema expressing cytosolic roGFP2-Orp1 (line #1, upper panel) or secreted $Ap1_{sp}$-roGFP2-Orp1 (line #131, middle and lower panels). Merge images indicate uniform sensor reduction in the cytosol. Sensor fluorescence was only visible in few protonema tip cells (middle panel), with uniform oxidation while most tip cells did not retain sensor signal in the apoplast. Images show full cell views of the cells depicted in Fig. 5. **B** In the transition from 2D to 3D growth in a you bud, sensor signal is visible in the apoplast of you leaflets, correlating with cuticle formation.

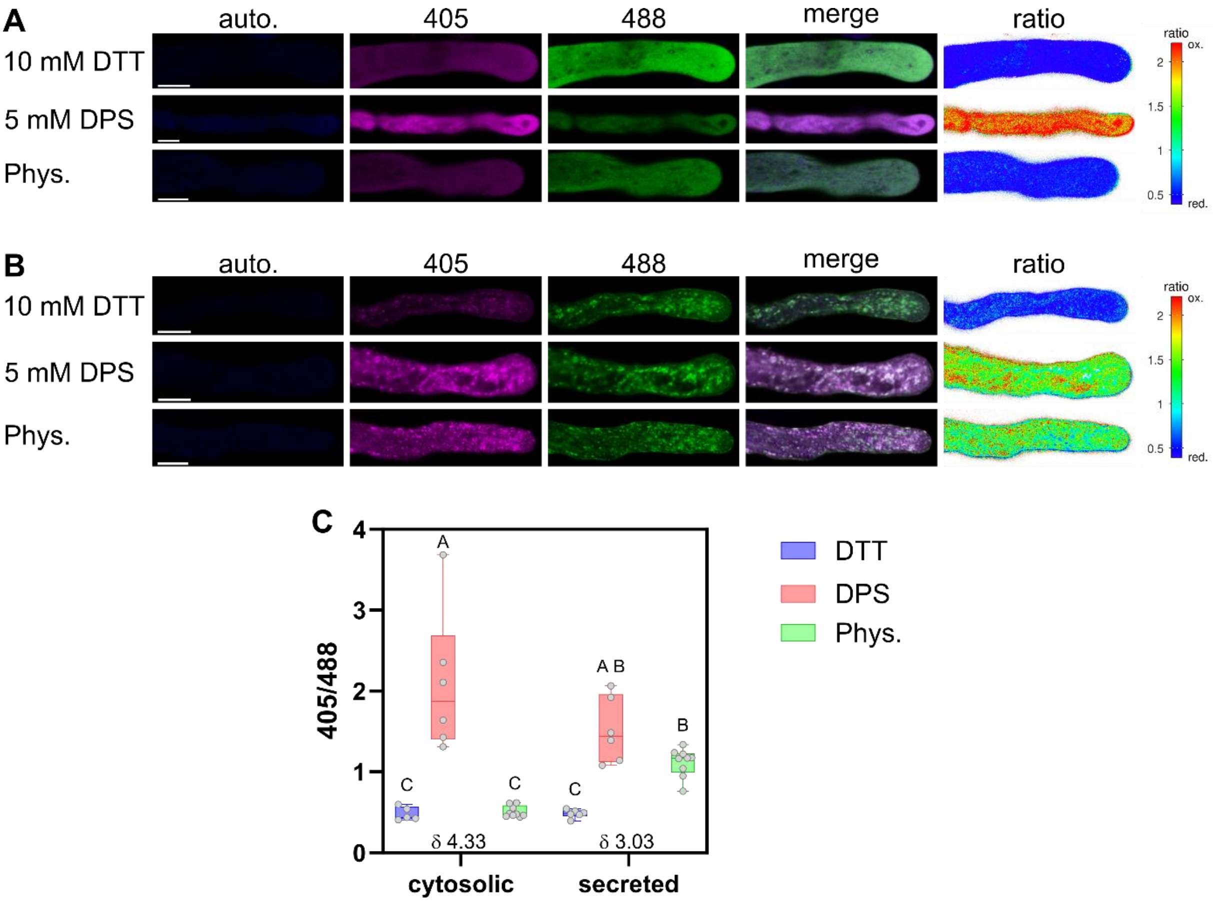


**Figure S13: Comparison of roGFP2-Orp1 in the cytosol and in the apoplast of *N. tabacum* pollen tubes.** **A**, **B** Exemplary confocal images showing *N. tabacum* pollen tubes expressing cytosolic (A) or secreted (B), *roGFP2-Orp1* or *CHI*$_{sp}$*-roGFP2-Orp1*, respectively. *roGFP2-Orp1* under fully reducing (DTT), fully oxidising (DPS) and physiological (phys.) conditions. Autofluorescence (auto.) elicited after excitation at 405 nm is depicted in blue (λem: 425 - 475 nm), roGFP2 signal after excitation at 405 nm in magenta (λem: 509 - 535 nm) and the roGFP2 signal after excitation at 488 nm in green (λem: 509 - 535 nm). The 405/488 fluorescence ratio is shown as merge of both roGFP signals (merge) and as false-colour-coded ratio (ratio); scale bars = 5 µm. **C** Box plots of ratio analysis using confocal microscopy images, measured dynamic range is indicated by δ. Boxes show 25th to 75th percentiles, line indicates the median, whiskers show min and max, individual data points are shown as circles, n = 5-9. Different letters indicate significant differences according to a two-way ANOVA with Tukey's multiple comparison post-hoc test, p<0.05.
